# Postnatal reduction of eIF4E overexpression in D1-SPNs ameliorates KCNQ dysfunction, hyperexcitability and ASD-like behaviours

**DOI:** 10.1101/2025.01.18.633694

**Authors:** Alina Aaltonen, Ayu Tamaki, Andrés Peris Ramón, Anders Borgkvist, Emanuela Santini

## Abstract

An imbalance between the direct and indirect pathways of the striatum has been implicated in the pathophysiology of ASD, which corresponds with an increase in repetitive behaviours and hyperactivity. The ASD risk gene *EIF4E* promotes translation, and its overexpression in mice increases repetitive behaviours and hyperactivity. We used the eIF4E-transgenic mouse model of ASD to study cell-type specific disruptions in the direct and indirect pathways using fibre photometry, electrophysiology, conditional gene silencing, and behavioural analysis. We found that direct pathway SPNs activity increased during exploratory behaviour and identified D1-SPN hyperexcitability and reduced KCNQ channel function in striatal slices. Reduction of eIF4E specifically in the D1-SPNs of adult mice normalised KCNQ function, D1-SPN hyperexcitability and ameliorated repetitive and hyperactive behaviours. Our results highlight the critical role of eIF4E in ASD-associated motor behaviours, elucidate cell-specific mechanisms driving hyperactivity and provide new insight into potential therapeutic targets for ASD and other neurodevelopmental disorders. Overall, this study underscores the translational potential of modulating protein synthesis pathways to address core motor symptoms in ASD.

## Introduction

Autism spectrum disorder (ASD) is a neurodevelopmental condition defined by core symptoms such as stereotyped, repetitive behaviours and deficits in social interactions and communication^1^. These behaviours are key for clinical diagnosis, yet the heterogeneity of ASD symptomatology and significant overlap with other neurodevelopmental and psychiatric conditions complicate diagnosis. This complexity underscores the need to better ascertain the neuronal circuits involved in ASD to develop effective biomarkers and targeted therapies. Recent MRI and fMRI studies have implicated specific brain regions in ASD pathology, including the striatum, the main input nucleus of the basal ganglia. Individuals with ASD show atypical striatal development^2–4^, altered striatal volume^5–7^, and disrupted corticostriatal connectivity^8,9^. In addition, preclinical investigations have further established that striatal dysfunctions contribute to ASD-like motor behaviours, such as stereotypies and perseverative behaviours^10–18^. Together, these findings suggest that striatal abnormalities play a critical role in the expression of ASD-related motor symptoms and repetitive behaviours^19–22^.

The striatum mainly consists of GABAergic spiny projection neurons (SPNs), which form two anatomically and functionally distinct striatal output pathways: the direct striatonigral and indirect striatopallidal pathways. These pathways, involving D1- and D2-SPNs, respectively, are central to movement regulation^23–25^. Although initially conceptualised as operating in an antagonistic balance^26–28^, recent evidence instead suggests that both pathways are coactivated during movement^25,29^. These findings support models where either direct-pathway SPNs drive specific motor programs and indirect-pathway SPNs suppress competing actions^25,29,30^, or coordinated activity between D1- and D2-SPNs within specific ensembles determines the motor program executed^31^. It has also been proposed that motor actions may result from a combination of these mechanisms^32^. Dysfunction in either pathway can disrupt motor control, resulting in disorders characterised by hyperactivity or stereotyped behaviours. Hyperactivity in neurodevelopmental and psychiatric disorders may result from inadequate suppression of actions by the indirect pathway^10,16,33^ or excessive activity in the direct pathway^13,15,34^. Therefore, identifying specific disruptions in D1- or D2-SPNs in neurodevelopmental disorders could reveal mechanisms underlying motor impairments and hyperactivity.

Aberrant protein synthesis has been implicated as a potential convergent mechanism across both genetic and idiopathic ASD-associated disorders^35–37^. Several ASD-implicated genes converge on the mammalian target of rapamycin (mTOR) signalling pathway^35,37^. The *EIF4E* gene encoding the cap-binding protein eIF4E, involved in mTOR-dependent translation, has been identified as an ASD risk gene in humans^38^. Dysregulated eIF4E function affects neurodevelopment^39^, and global^40,41^ or microglia-specific overexpression of eIF4E^42^ or knockout of *4E-BP2* encoding the eIF4E repressor protein^43^ results in aberrant synaptic function and neuronal excitability as well as ASD-like behaviours, including increased grooming, marble burying and novelty-induced hyperlocomotion^40–43^. Thus, examining the effect of eIF4E overexpression may provide insight into the mechanism by which specific disruptions in SPNs contribute to ASD-related behaviours.

In this study, we utilised the eIF4E-transgenic (eIF4E-TG) mice to test the hypothesis that altered excitability in the D1- or D2-SPNs causes an imbalance in the striatonigral and striatopallidal pathways that drives hyperactivity and repetitive behaviours. By integrating *in vivo* neuronal activity measurements, *ex vivo* patch clamp electrophysiology, and conditional gene silencing, we demonstrate that eIF4E overexpression leads to elevated Ca²⁺ activity in D1-SPNs during exploratory activity in freely behaving mice. Furthermore, we found that eIF4E-TG mice exhibit D1-SPN hyperexcitability after postnatal maturation and reduced conductance through voltage-gated Kv7 (KCNQ) potassium channels, which play a pivotal role in the control of membrane excitability. Importantly, reducing eIF4E overexpression in D1- SPNs in adult eIF4E-TG mice increased the KCNQ current, reduced hyperexcitability, and normalised novelty-induced motor behaviours. These findings underscore the importance of understanding cell-specific mechanisms, as targeting D1-SPNs may offer new therapeutic approaches for motor dysfunctions in ASD and related disorders.

## Results

### Increased novelty-induced striatonigral activity in the eIF4E-TG mice

The eIF4E*-*transgenic (eIF4E-TG) mice exhibit repetitive behaviours and hyperactivity when introduced to a novel, unfamiliar environment^40^, which stimulates concurrent direct and indirect pathway activity associated with locomotion^29,44^. To explore whether dysfunctions in the striatal pathways are responsible for the hyperlocomotion observed in the eIF4E-TG mice, we assessed novelty-induced changes in fluorescence of the genetically encoded Ca^2+^ indicator GCaMP7s used as a proxy for spiking activity in direct and indirect SPNs using fibre photometry. We injected Cre-inducible GCaMP7s into the dorsal striatum (Fig. 1a) of mice generated by crossing the eIF4E-TG line with D1- and A2a-Cre mice (D1-Cre/WT and -/TG and A2a-Cre/WT and -/TG; see materials and methods).

**Figure 1:**
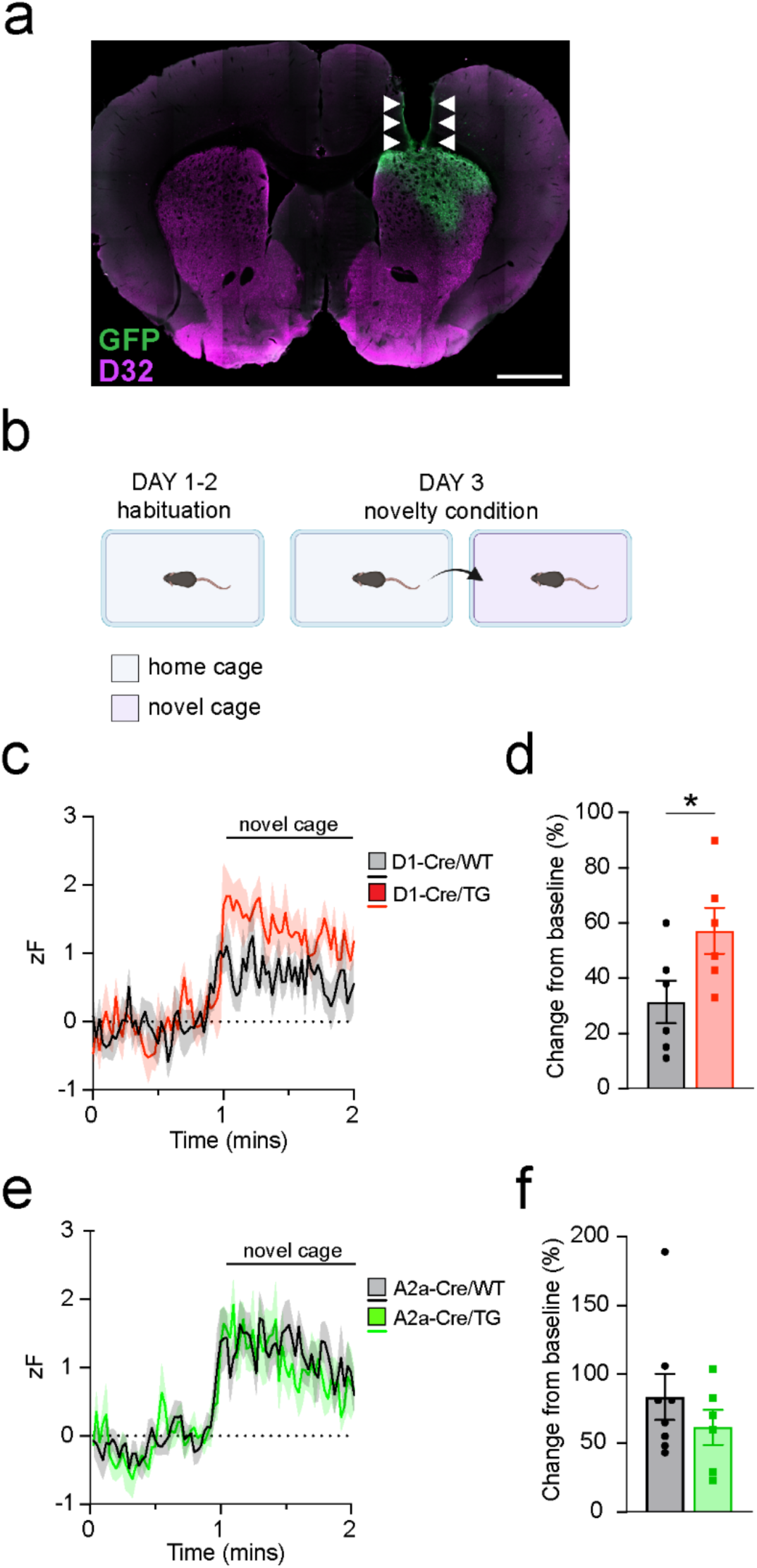
Novelty reveals increased activity in striatonigral SPNs of eIF4E-TG mice. **a**) Coronal brain slice showing example of electrode placement within the dorsal striatum, with GCaMP7s expression shown in green, and DARPP32 immunostaining (indicating the striatum) in magenta. Scale bar indicates 1mm. **b**) Schematic of the experimental protocol, with two days of habituation to an empty home cage (blue) followed by one day of recording, including initial habituation to home cage (blue) with patch cable, followed by baseline recording before transfer to novel environment (pink). **c**) Averaged Z-score normalised GCaMP7s traces in D1-Cre/WT and -/TG mice (6 WT and 6 TG). Z-scores are calculated from the pre-novelty baseline activity. **d**) Change in GCaMP7s activity, shown as a combined average of three distinct parameters relating to activity (Supplemental Fig. 2a, c and e>), expressed as a change from baseline, for the direct pathway SPNs (unpaired T-test: t=2.27, *p=0.0465). **e**) Averaged Z-score normalised GCaMP7s traces, during the timepoint one minute before until one minute after introduction to novel environment, in A2a-Cre/WT and -/TG mice (8 WT and 6 TG). Z-scores are calculated from the pre-novelty baseline activity. **f**) Change in GCaMP7s activity, shown as a combined average of three distinct parameters relating to activity (Supplemental Fig. 2b, d and f>), expressed as a change from baseline, for the indirect pathway SPNs. In all panels, data are presented as mean ± SEM, with dots representing individual values for each mouse. The mean is indicated by the bar height or connecting line, while the SEM is shown by error bars or lines with shading. For full details of statistical analysis including negative results, refer to supplemental table 1.

We recorded baseline GCaMP7s fluorescence first in a familiar home cage and then in a novel environment (Fig. 1b). To assess the novelty response, GCaMP7s fluorescence after novel cage exposure in each mouse was normalised to its baseline, yielding a percentage change in fluorescence. Exposure to the novel environment enhanced the Ca^2+^ activity in both direct and indirect SPNs in WT mice (Fig. 1c,e) (D1: 31.33% WT, one-sample T-test: t(5)=4.057, p=0.0098; A2a: 83.5% WT, one-sample T-test: t(7)=4.986, p=0.0016). Importantly, the eIF4E- TG mice exhibited an enhanced novelty-induced GCaMP7s fluorescence in direct SPNs compared to WT, but not in indirect SPNs (Fig. 1c,d and 1e,f). Further analysis of individual parameters revealed increases in peak frequency (Supplemental Fig. 1a) and area under the curve (Supplemental Fig. 1e) in direct SPNs of eIF4E-TG mice, with no change in signal amplitude (Supplemental Fig. 1c). In indirect SPNs, no difference between eIF4E-TG mice and WT was observed in any parameter (Supplemental Fig. 1b, d, f>).

To ensure that our *in vivo* measurements reported the correct cell-type specific changes in Ca^2+^ signal, we performed post hoc tissue analysis with immunostaining using antibodies detecting GCaMP7s and DARPP-32, a marker for SPNs. We confirmed a 50% co-expression of GCaMP7s and DARPP-32 in the respective experimental groups (Supplemental fig. 2a-b), in agreement with the equinumerous distribution of D1- and D2-SPNs^45^. In addition, treatment of D1-Cre/WT mice with the D1-receptor selective antagonist SCH39166 reduced D1-SPN GCaMP7s fluorescence (Supplemental Fig. 2d-g>). Fibre photometry recordings performed on mice expressing GFP did not reveal any movement-related changes in the parameters we used to quantify changes in GCaMP7s fluorescence^29^ (Supplemental Fig. 2c). Altogether, our results show that the eIF4E-TG mice exhibit a direct pathway-specific hyperactivity in response to novelty.

### Overexpression of eIF4E results in cell-type specific hyperexcitability in SPNs

Our fibre photometry recordings suggested that eIF4E-TG mice exhibit electrophysiological changes in SPNs reminiscent of FXS mouse models, which share molecular synaptic and behavioural phenotypes with the eIF4E-TG mice^15,46^. Therefore, we investigated whether alteration in membrane conductance maintaining excitability might underlie the hyperactivity observed in D1-SPNs *in vivo*. We performed whole-cell patch clamp electrophysiology in striatal slices obtained from Drd1a-TdTomato mice^47^ crossed with the eIF4E-TG line (D1- Tomato/WT and -/TG mice) to isolate TdTomato-fluorescent D1-positive or D1-negative SPNs (the latter presumed to be D2-SPNs). In D1-SPNs, we found no significant differences between TG and WT littermates in membrane capacitance (Fig. 2a) or resting membrane potential (RMP, Fig. 2b). However, membrane resistance of D1-SPNs was significantly increased in eIF4E-TG mice compared to WT mice (Fig. 2c), indicating genotype-related differences in voltage-gated channel composition, conductance or density.

**Figure 2:**
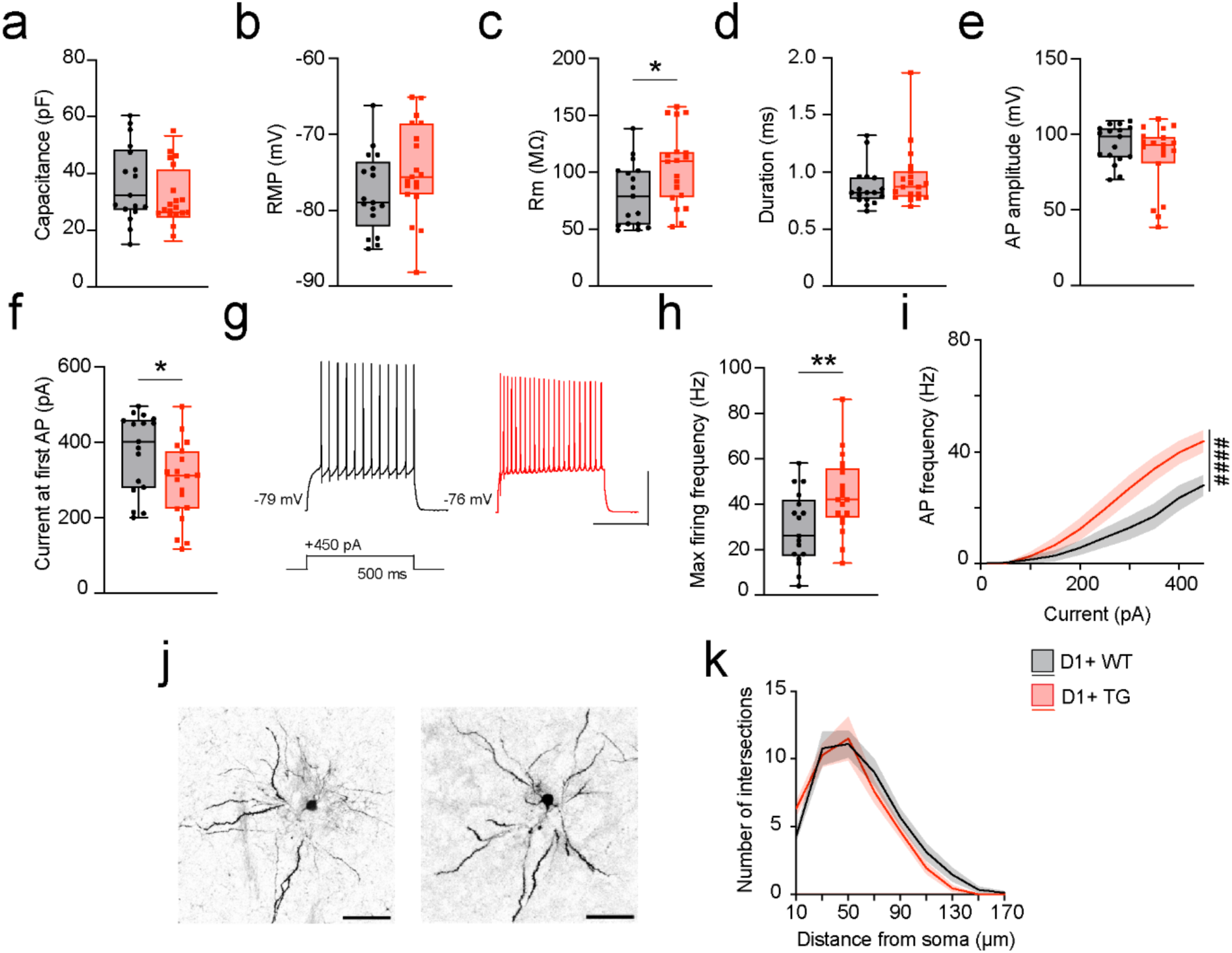
Increased excitability in D1-SPNs of eIF4E-TG mice **a**) Calculated capacitance in D1-SPNs from D1-Tomato/WT and D1-Tomato/TG. **b**) Resting membrane potential. **c**) Membrane resistance (Rm, unpaired T-test: t=2.267, *p=0.0299). **d**) Action potential half-height duration. **e**) Action potential peak amplitude. **f**) Rheobase current (unpaired T-test: t=2.267, *p=0.0298). **g**) Representative traces of action potentials at +450pA current injection for 500ms (scale bar: 200ms, 40mV). **h**) Maximum firing frequency across the range of injected currents (unpaired T-test: t=2.765, **p=0.0091). **i**) Current-frequency (IF) plots for D1-SPNs (currentXgenotype interaction, RM-2-way ANOVA: F(9, 306)=5.547, #### p<0.0001). **j**) Representative images of Lucifer Yellow-filled D1-SPNs (scale bar: 50µm). **k**) Dendritic intersections plotted as a function of distance from soma. Data are presented as median ± quartiles within the box plot, with whiskers indicating minimum and maximum values (**a**-**h**), or as mean ± SEM, with the mean represented by a connecting line and SEM by error lines with shading (**i** and **k**). Dots represent individual values for each neuron. All recordings were collected from D1+ cells (n=17 WT, n=19 TG) from 5 D1-Tomato/WT and 5 D1- Tomato/TG mice. Significance is denoted as * p<0.05, ** p<0.01, *** p<0.001 and **** p<0.0001, calculated using Student’s t-test. For full details of statistical analysis, including negative results, refer to Supplemental table 2.

We did not observe differences in action potential (AP) kinetics such as AP half-height duration (Fig. 2d) or AP peak height (Fig. 2e). However, the rheobase current in D1-SPNs was reduced (Fig. 2f), accompanied by increases in maximum firing frequency (Fig. 2g-h) and current-frequency responses (Fig. 2i), indicating a lower threshold for firing and heightened excitability. Since differences in membrane properties could reflect morphological changes^48^, we performed Sholl analysis on D1-SPNs microinjected with the fluorescent dye Lucifer Yellow. However, no difference was found in the distribution of dendritic intersections or cell size (Fig. 2 j-k). These findings show that D1-SPNs of the eIF4E-TG mice exhibit normal dendritic morphology and unaltered RMP, but are hyperexcitable during depolarisation, which could contribute to the hyperactivity phenotype observed *in vivo*.

We also analysed the intrinsic properties of D2-SPNs (Supplemental Fig. 3) and found a reduction in capacitance in eIF4E-TG mice (Supplemental Fig. 3a), indicating that there may be differences in cell size in D2-SPNs. However, we did not observe a change in the dendritic morphology of D2-SPNs (Supplemental Fig. 3j-k>). In summary, we identified increased excitability specifically in D1-SPNs of eIF4E-TG mice, as evidenced by a reduced rheobase and enhanced firing response to depolarising current injection. Given the absence of morphological or structural changes, the observed increase in membrane resistance in D1- SPNs likely reflects alterations in membrane excitability, potentially due to changes in voltage-gated ion channel composition, conductance or density.

### D1-SPN hyperexcitability develops after SPN maturation

SPNs undergo extensive maturation of their electrophysiological and structural properties between the first and fourth postnatal weeks, ultimately gaining adult-like phenotypes^49–51^. Because eIF4E is overexpressed prenatally in eIF4E-TG mice, we sought to determine whether the differences in neuronal properties of D1-SPNs emerge during a particular phase of postnatal development. To test this, we recorded the intrinsic properties and depolarisation-induced membrane responses of D1 and D2-SPNs at postnatal day P10 (early in the developmental period), and P28 (towards the end of the period)^52^, comparing D1-Tomato/WT and D1-Tomato/TG mice.

While the capacitance of D1-SPNs did not change during this interval (Fig. 3a), we found a significant age-associated decrease in RMP (Fig. 3b) and membrane resistance in WT mice (Fig. 3c), as previously reported^52^. Interestingly, D1-SPNs in eIF4E-TG mice exhibited mature-like RMP at P10 (Fig. 3b) and a lack of reduction in membrane resistance between P10 and P28 (Fig. 3c), indicating that genotype-related differences in membrane properties are established postnatally. No maturation-dependent differences were found in AP half-height duration (Fig. 3d) or AP peak amplitude (Fig. 3e). We also did not observe any significant genotype differences in the age-associated increase in rheobase current or the reduction in maximum firing frequency (Fig. 3f-h), confirming that D1-SPN excitability decreases in both eIF4E-TG and WT mice during postnatal development. However, we found that D1-SPNs of eIF4E-TG mice responded to varying current injections with more action potentials at P28 but not at P10 (Fig. 3g, i). In contrast, D2-SPNs showed no genotype-related differences in any of the parameters measured (Supplemental Fig. 4). In conclusion, we observed dysregulation in the maturation of passive membrane properties in D1-SPNs but little alteration in the maturation of excitability during postnatal development in the eIF4E-TG mice. These findings suggest that the differences in D1-SPN excitability and firing patterns emerge after P28 when SPNs have gained adult-like membrane properties.

**Figure 3:**
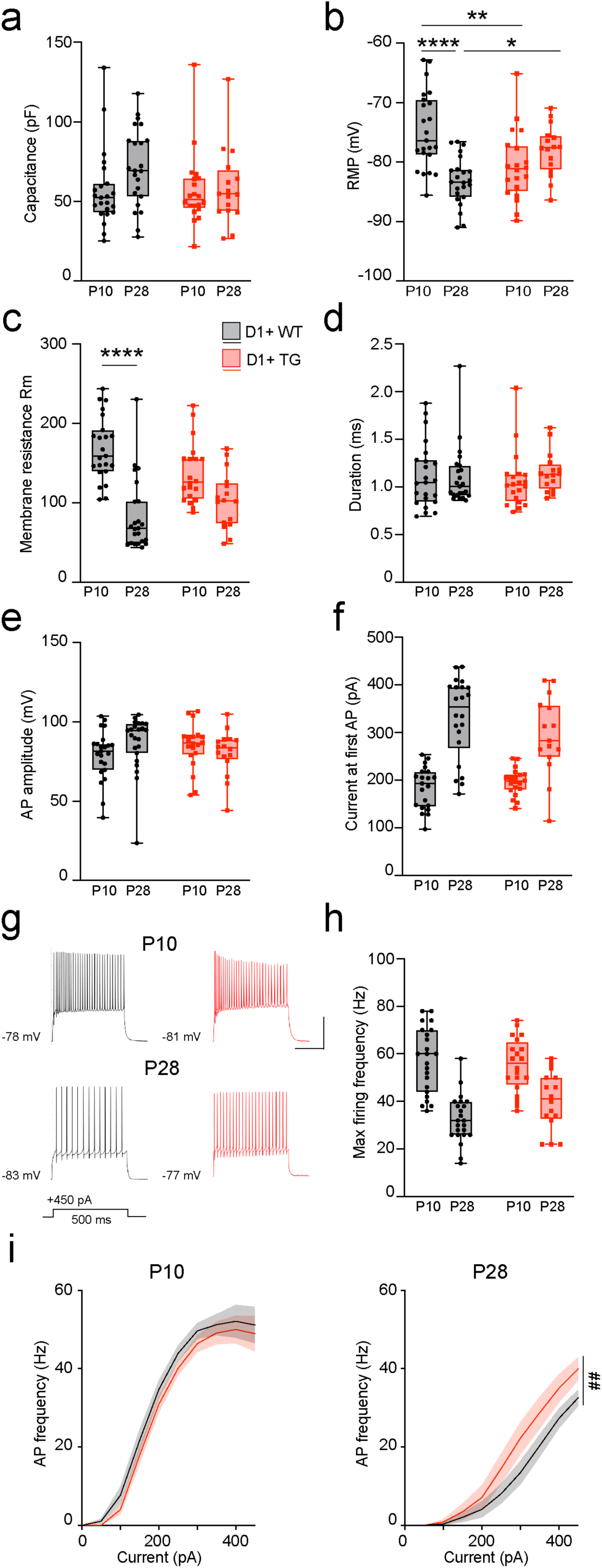
D1-SPN hyperexcitability emerges after the postnatal developmental period **a**) Calculated capacitance of D1-SPNs from D1-Tomato/WT and D1-Tomato/TG at P10 and P28. **b**) Resting membrane potential in D1-SPNs of P10 and P28 mice (genotypeXage interaction, 2-way ANOVA: F(1, 77)=21.06, ****p<0.0001). **c**) Membrane resistance of D1- SPNs at P10 and P28 (genotypeXage interaction, 2-way ANOVA: F(1, 78)=7.49, **p=0.0077). **d**) Action potential half-height duration of D1-SPNs at P10 and P28. **e**) Action potential peak amplitude of D1-SPNs at P10 and P28. **f**) Rheobase current of D1-SPNs at P10 and P28. **g**) Representative traces of action potentials at +450pA current injection for 500ms (scale bar: 200ms, 50mV). **h**) Maximum firing frequency across the range of injected currents in D1-SPNs at P10 and P28 mice. **i**) Current-frequency (IF) plots for D1-SPNs at P10 and P28 (P28: currentXgenotype interaction, repeated measures 2-way ANOVA: F(9, 333)=2.635, ##p=0.0059). Data are presented as median ± quartiles within the box plots, with whiskers indicating minimum and maximum values (**a**-**h**), or as mean ± SEM, with the mean represented by a connecting line and SEM by error lines with shading (**i**). Dots represent individual values for each neuron. Recordings were collected from D1+ cells from 7 D1-Tomato/WT and 5 D1- Tomato/TG P10 mice and 5 D1-Tomato/WT and 4 D1-Tomato/TG P28 mice (n=23 WT P10, n=20 TG P10; n=23 WT P28, n=16 TG P28). Significance is denoted as * p<0.05, ** p<0.01, *** p<0.001 and **** p<0.0001, calculated using Tukey’s post-hoc multiple comparisons test. For full details of statistical analysis including negative results, refer to supplemental table 3.

### Conditional silencing of eIF4E reduces firing frequency of D1-SPNs

Given the cell-type specificity and postnatal establishment of the firing abnormalities observed in eIF4E-TG mice, we hypothesised that the dysregulation occurs in a cell-autonomous manner and that reducing eIF4E levels in D1-SPNs in adulthood could reverse the impairments. To reduce eIF4E expression in adult mice, we utilised an inducible RNAi approach similar to that previously established with *Eif4e*-specific shRNA (shmiR-4E) constructs^53–55^. This approach incorporates a knock-in, Tet-on-based conditional expression system, where shmiR-4E is embedded within the *Gfp* sequence and regulated by tet-responsive elements (TRE), enabling a conditional reduction of eIF4E protein levels^53,55^.

We assessed the efficacy of the double-conditional gene expression system for reducing eIF4E in SPNs in the striatum of TRE-GFP.shmiR-4E mice. We co-infused an AAV-Cre together with an AAV-DIO-tTA transcribing a neuron-specific Cre-dependent tet-transactivator (tTA) into the striatum. Cre-induced activation of tTA will express shmiR-4E in infected neurons, resulting in eIF4E reduction and concomitant GFP expression marking the targeted cells. As control, we co-injected AAV-*Cre* with an AAV-FLEX-TdTomato to visualise double-infected control cells. Three weeks post-surgery, we observed a reduction of eIF4E levels in GFP-positive neurons of tTA-injected mice (Supplemental Fig. 5a) but not in TdTomato-positive control neurons (Supplemental Fig. 5b) of TdTomato-injected mice. Furthermore, we found a significant decrease in eIF4E-positive neurons in the striatum of tTA-injected mice compared to those of TdTomato-injected mice (Supplemental Fig. 5c). Both GFP and TdTomato were predominantly expressed in DARPP-32-positive neurons, with 79% of fluorophore-positive neurons (i.e. Cre and either Cre-inducible tTA or Cre-inducible TdTomato) containing DARPP-32 (Supplemental Fig. 5d-e>). To quantify eIF4E reduction, we performed Western Blot analysis on striatal tissue extracts and confirmed a substantial reduction in eIF4E in samples from AAVs-injected double transgenic TRE-GFP.shmiR-4E/ WT and -/TG littermates, with no significant genotype differences (Supplemental Fig. 5f-g>).

To reduce eIF4E expression specifically in D1-SPNs, we injected a retrograde AAV-Cre into the substantia nigra pars reticulata (SNr) of TRE-GFP.shmiR-4E/ WT mice, selectively targeting direct pathway SPNs, along with AAV-DIO-tTA in the striatum in the left hemisphere (tTA^L^), and AAV-FLEX-TdTomato in the right hemisphere (TdTomato^R^) (Fig. 4a). Approximately 95% of striatal neurons are SPNs—with roughly equal distribution of D1- and D2-SPNs^45^, which can be revealed by immunostaining with antibodies against enkephalin that is expressed selectively by D2-SPNs^56,57^. In striatal brain sections from double-injected mice, we observed that 93% of fluorophore-positive cells were enkephalin-negative (Supplemental Fig. 5h-i>), confirming that our approach selectively targets D1-SPNs.

**Figure 4:**
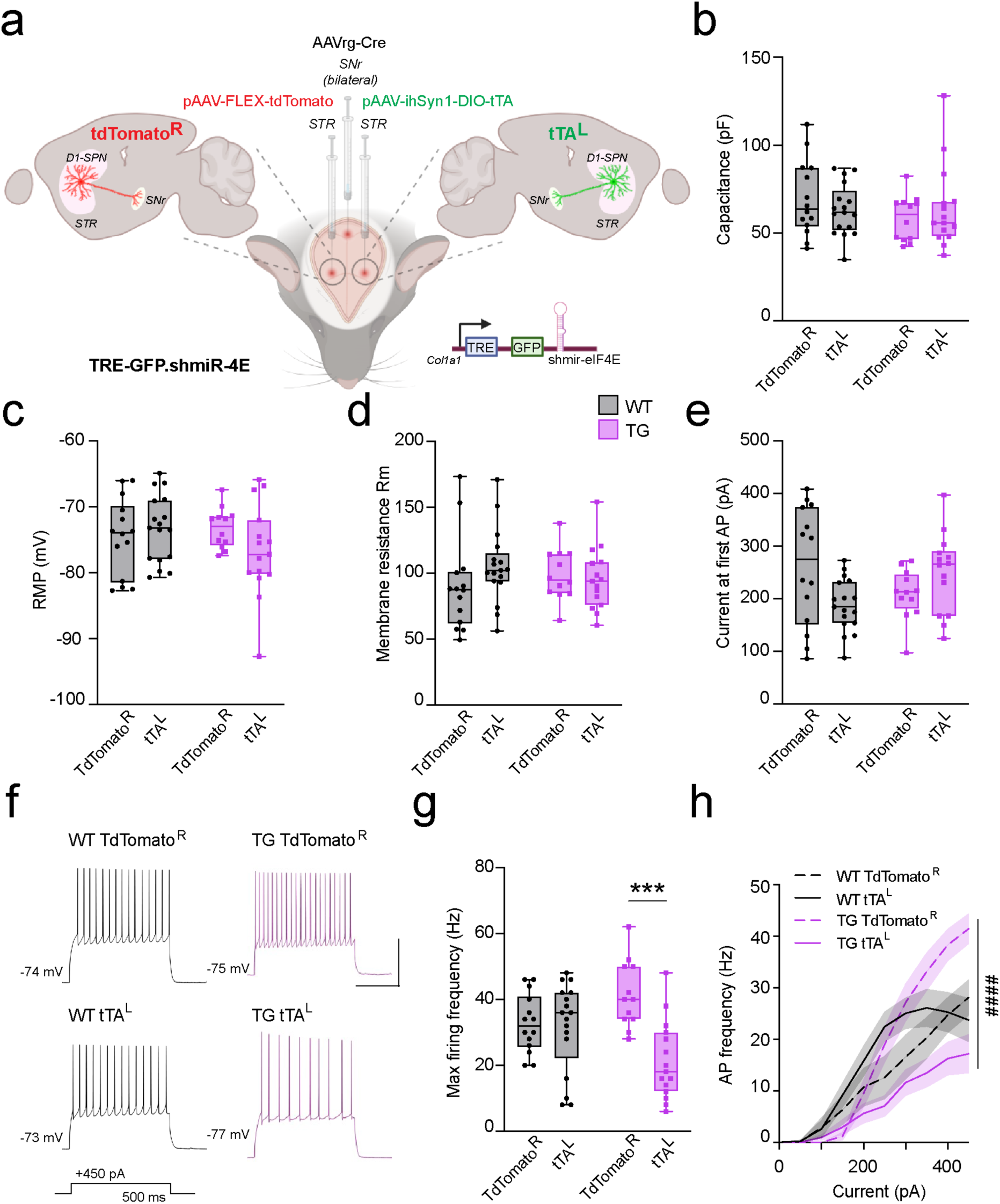
Conditional reduction of eIF4E in adult D1-SPNs decreases firing frequency **a**) Schematic of the experimental design to selectively reduce eIF4E D1-SPNs. A retrograde AAVrg-*Cre* was injected into the substantia nigra pars reticulata (SNr) of TRE-GFP.shmiR-4E/ WT and -/TG mice to target D1-SPNs. An AAV-ihSyn1-DIO-tTA was injected into the left striatum (tTA^L^), activating RNAi for eIF4E and driving GFP expression in D1-SPNs. In parallel, an AAV-FLEX-tdTomato was injected into the right striatum (TdTomato^R^), driving the expression of a control fluorescent marker in D1-SPNs. **b**) Capacitance in D1-SPNs from TRE-GFP.shmiR-4E/ WT and -/TG mice in tTA^L^ or TdTomato^R^ hemispheres. **c**) Resting membrane potential in D1-SPNs from tTA^L^ or TdTomato^R^ hemispheres. **d**) Membrane resistance in D1- SPNs from tTA^L^ or TdTomato^R^ hemispheres. **e**) Rheobase current in D1-SPNs from tTA^L^ or TdTomato^R^ hemispheres (genotypeXRNAi interaction, 2-way ANOVA: F(1, 53)=8.028, **p=0.0065). **f**) Representative traces of action potentials at +450pA current injection for 500ms (scale bar: 200ms, 50mV). **g**) Maximum firing frequency D1-SPNs from tTA^L^ or TdTomato^R^ hemispheres (genotypeXRNAi interaction, 2-way ANOVA: F(1, 54)=10.10, **p=0.0024). **h**) Current-frequency (IF) plots for D1-SPNs from tTA^L^ or TdTomato^R^ hemispheres (currentXgenotypeXRNAi interaction, 3-way ANOVA: F(9, 486)=6.46, #### p<0.0001). Data are presented as median ± quartiles within the box plots, with whiskers indicating minimum and maximum values (**a**-**g**), or as mean ± SEM, with the mean represented by a connecting line and SEM by error lines with shading (**h**). Dots represent individual values for each neuron. All recordings were collected from 4 TRE-GFP.shmiR-4E/ WT and 4 TRE-GFP.shmiR-4E/ TG mice (n=14 WT TdTomato^R^, n=17 WT tTA^L^, n=12 TG TdTomato^R^, n=15 TG tTA^L^). Significance is denoted as * p<0.05, ** p<0.01, *** p<0.001 and **** p<0.0001, calculated with Tukey’s post-hoc multiple comparisons test. For full details of statistical analysis including negative results, refer to supplemental table 4.

We then recorded intrinsic properties of D1-SPNs in striatal brain slices from TRE-GFP.shmiR- 4E/ WT and -/TG mice (Fig. 4a). The experimental design enabled us to compare the properties of eIF4E-reduced tTA^L^ D1-SPNs and control TdTomato^R^ neurons from the same mouse. Reducing eIF4E did not change the capacitance, RMP or membrane resistance of the D1-SPNs (Fig. 4b-d). Importantly, however, we found an interaction effect in the rheobase current (Fig. 4e), and a reversal of the firing frequency dysregulation observed in eIF4E-TG mice (Fig. 4f-h). Thus, reducing eIF4E levels in adult D1-SPNs of eIF4E-TG mice normalised the aberrant firing activity caused by global eIF4E overexpression.

### Potassium channelopathies in D1-SPNs

An increasing body of evidence links various channelopathies to ASDs, including those of K+ channels^58^. SPNs highly express Kir2 channels, which maintain the hyperpolarised resting membrane potential observed in mature SPNs^52,56,59^. Although the RMP of D1-SPNs was similar in adult eIF4E-TG and WT mice, D1-SPNs in eIF4E-TG exhibited higher membrane resistance, which could result from alterations in Kir2 channel activity^52^. To exclude this possibility, we subjected D1-SPNs from D1-Tomato/WT and -/TG mice to hyperpolarising voltage steps within the voltage-range of Kir2 channel activation and then applied the K- channel blocker Cs+ to extract the Kir2 channel current^52^. We found no difference in the amplitude of the Cs+-sensitive current between eIF4E-TG and WT (Fig. 5a-b), suggesting that Kir2 channel dysfunction is not responsible for the D1-SPN hyperexcitability observed in eIF4E-TG mice.

**Figure 5:**
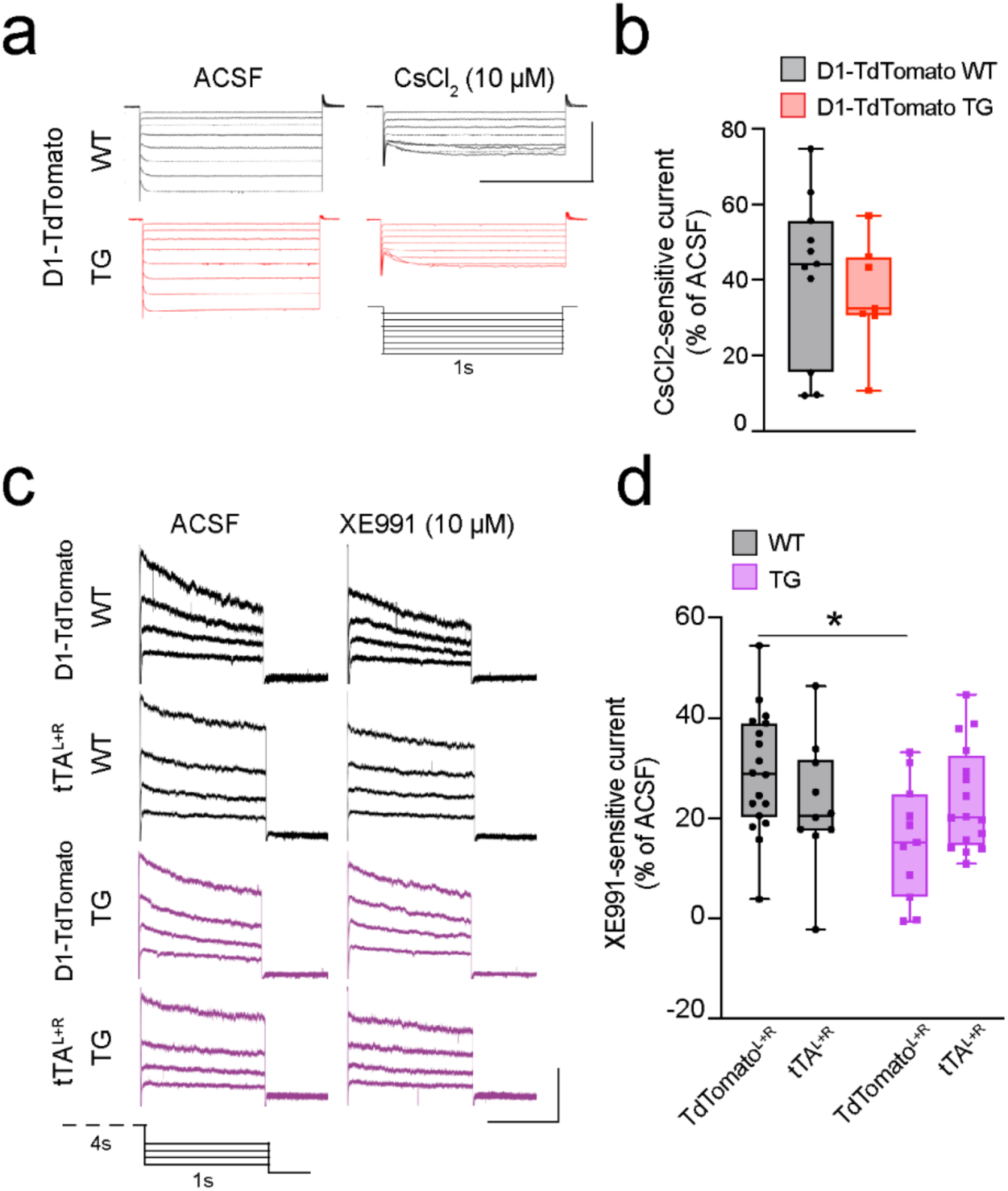
Reduced KCNQ function in D1-SPNs of the eIF4E-TG mice is normalised by conditional eIF4E reduction **a**-**b**) Recordings conducted in the presence of CsCl2 were collected from D1+ cells (n=11 WT, n=7 TG) from 6 D1-Tomato/WT and 4 D1-Tomato/TG mice. **a**) Representative traces from D1- Tomato/WT and -/TG D1-SPNs before (ACSF) and after (CsCl2) bath application of CsCl2, with one-second current steps from holding potential of -70mV to -140mV in 10mV increments (scale bar: 600ms and 300pA). **b**) Comparison of the percentage current reduction by CsCl2 at the most hyperpolarising voltage step between WT and TG D1-SPNs. **c**-**d**) Recordings with XE991 in ACSF were collected from D1+ cells (n=18 D1-Tomato/WT, n=11 D1-Tomato/TG) from 5 D1-Tomato/WT and 5 D1-Tomato/TG mice, and from D1+ cells (n=10 tTA^L+R^ WT and n=16 tTA^L+R^ TG) from 4 TRE-GFP.shmiR-4E/WT tTA^L+R^ and 6 TRE-GFP.shmiR-4E /TG tTA^L+R^ mice. **c**) Representative traces from D1-SPNs from D1-Tomato/ WT, -/ TG, TRE-GFP.shmiR-4E/WT tTA^L+R^ (WT tTA^L+R^ ) and -/TG tTA^L+R^ (TG tTA^L+R^ ) before (ACSF) and after (XE991) bath application of XE991, with one-second current steps from a brief (four second) holding potential of 0mV and from -20 to -50mV in 10mV increments (scale bar: 600ms and 300pA). **d**) Percentage current reduction by XE991 -20mV, comparing D1-Tomato/WT, -/TG, GFP.shmiR-4E/WT tTA^L+R^ and -/TG tTA^L+R^ D1-SPNs (genotypeXRNAi interaction, 2-way ANOVA: F(1, 51)=5.218, *p=0.0266; * p<0.05 Tukey’s post-hoc multiple comparisons test). All data are presented as median ± quartiles within the box plot, and whiskers indicate minimum and maximum values. Dots represent individual values for each neuron. For full details of statistical analysis including negative results, refer to supplemental table 5.

SPNs express multiple K+ channel subtypes^59^, including voltage-gated KCNQ channels that specifically sustain a non-activating K+ current (M-current) that reduces D1-SPN excitability^60,61^. Therefore, we examined whether impaired KCNQ channel function contributes to the observed D1-SPN hyperexcitability in eIF4E TG mice. We performed patch clamp recordings in slices from D1-Tomato/WT and D1-Tomato/TG mice, and TRE-GFP.shmiR- 4E/WT and -/TG mice with reduced eIF4E levels from bilateral injections of AAVrg-Cre in the SNr and AAV-DIO-tTA in the striatum (tTA^L+R^). To isolate the M-current, we subjected D1-SPNs to KCNQ-activating voltage-step commands first in standard ACSF and then in the presence of the KCNQ2/3 blocker XE991^60,62^. Supporting altered KCNQ function in D1-SPNs of the eIF4E-TG mice, we discovered a significantly smaller XE991-sensitive current in D1- TdTomato/TG mice compared to D1-TdTomato/WT (Fig. 5c-d). Strikingly, we found no significant difference between XE991-sensitive currents in D1-SPNs of TRE-GFP.shmiR- 4E/TG tTA^L+R^ mice compared to WT mice (Fig. 5c-d), confirming that reduced expression of eIF4E in D1-SPNs establishes normal KCNQ function. These findings suggest that the hyperexcitability of D1-SPNs in eIF4E-TG mice is mediated by KCNQ2/3 channel impairments, and that this effect is reversible through targeted reduction of eIF4E in adulthood.

### Reduction of eIF4E in D1-SPNs normalises hyperactivity and repetitive behaviours

As the reduction of eIF4E in D1-SPNs reduced hyperexcitability and increased KCNQ channel activity in adult mice, we hypothesised that reducing eIF4E levels in adult D1-SPNs decreases novelty-induced hyperlocomotion and ameliorates repetitive behaviours. To test this, we used TRE-GFP.shmiR-4E/WT and -TG littermates, injected with AAVrg-Cre in the SNr and either DIO-tTA (tTA^L+R^) or AAV-FLEXtdTomato (TdTomato^L+R^) bilaterally in the striatum, to achieve reduction of eIF4E in D1-SPN or expression of a control fluorophore. At least three weeks post-surgery, we first assessed novelty-induced locomotion as in the fibre photometry measurements. Subsequently, we performed the marble burying test, a paradigm previously used to quantify repetitive behaviour prevalent in ASD models^15,63^, including eIF4E-TG mice^40^.

Mice in all experimental groups habituated to the novel environment, showing a gradual decrease in the distance travelled over the course of two hours (Fig. 6a). However, TRE-GFP.shmiR-4E/TG mice expressing TdTomato^L+R^, which maintain elevated eIF4E in D1-SPNs, performed a significantly larger cumulative distance travelled compared to the tTA^L+R^ group, in which eIF4E levels were reduced (Fig. 6b). This difference could primarily be explained by an increased velocity (measured in cm/s averaged over five-minute time intervals, Supplemental Fig. 6a-b>), as the total movement duration (Supplemental Fig. 6c-d>) and immobile time (Supplemental Fig. 6e-f>) were unaffected by eIF4E reduction. Furthermore, inhibition of eIF4E overexpression in TRE-GFP.shmiR-4E/TG mice expressing tTA^L+R^ decreased the number of marbles buried to an indistinguishable level from WT mice injected with AAVs transducing TdTomato (Fig. 6c).

**Figure 6:**
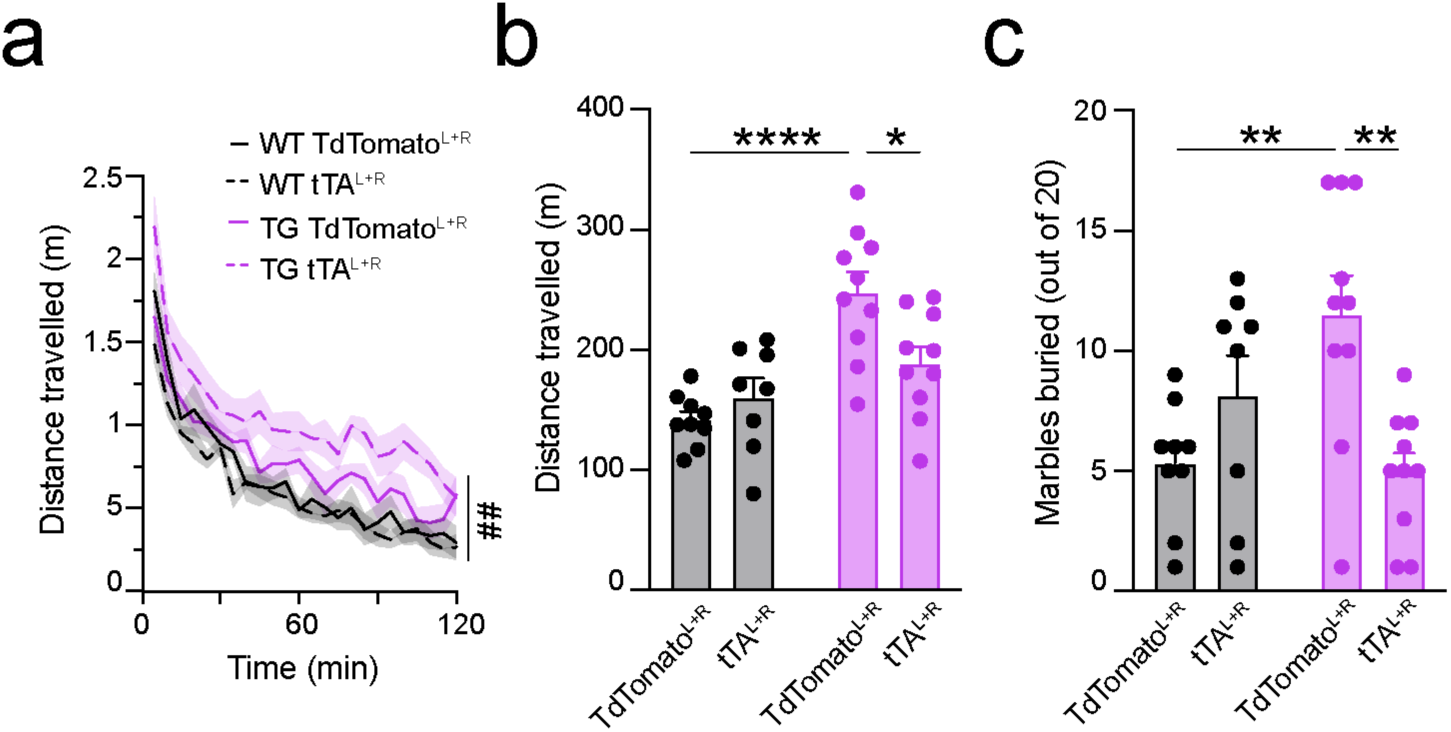
Conditional reduction of eIF4E in adult D1-SPNs ameliorates hyperactivity and repetitive behaviours in eIF4E-TG mice **a**) Time-course of the distance travelled during the novelty condition experiment, with data binned in five-minute intervals. **b**) Total distance travelled over the two-hour experimental period (genotypeXRNAi interaction, 2-way ANOVA: F(1, 33)=7.607, ## p=0.0094). **c**) Total number of marbles buried during the 30-minute marble burying experiment (genotypeXRNAi interaction, 2-way ANOVA, F(1, 33)=13.13, *p=0.018). Data are presented as mean ± SEM, with mean indicated by bar height or connecting line and SEM by error bars or lines with shading. Dots represent individual values for each neuron. Behavioural experiments were performed using 9 TRE-GFP.shmiR-4E/WT TdTomato^L+R^ and 10 TRE-GFP.shmiR-4E/TG TdTomato^L+R^ mice, and 8 TRE-GFP.shmiR-4E/WT tTA^L+R^ and 10 TRE-GFP.shmiR-4E/TG tTA^L+R^ mice. Significance is denoted as * p<0.05, ** p<0.01, *** p<0.001 and **** p<0.0001, calculated with Tukey’s post-hoc multiple comparisons test. For full details of statistical analysis including negative results, refer to supplemental table 6.

These findings demonstrate that eIF4E overexpression enhances D1-SPN excitability and striatonigral pathway activity *in vivo*, driving hyperactivity and repetitive behaviours. D1-SPN-targeted reduction of eIF4E overexpression in adulthood mitigates these ASD-related behaviours, highlighting a promising therapeutic strategy for addressing hyperactivity and repetitive behaviours in ASD.

## Discussion

Using whole-cell *ex vivo* patch clamp electrophysiology and *in vivo* fibre photometry, we have demonstrated that eIF4E overexpression, which alters global protein synthesis, results in cell type-specific functional dysregulation in the striatum. Our results show that D1-SPNs are more active during novelty-induced locomotion in the eIF4E-TG mice, and the D1-SPNs selectively exhibit altered intrinsic properties suggestive of hyperexcitability and hyperactivity, which develop during the postnatal developmental period. Additionally, selective reduction of eIF4E in adult D1-SPNs is sufficient to normalise enhanced firing frequency, impaired KCNQ channel activity and ASD-like behaviours such as novelty-induced hyperlocomotion and marble burying. This suggests that altered D1-SPN activity driven by eIF4E overexpression underlies these ASD-related cellular and behavioural phenotypes.

Exploratory behaviour in novel environments relies on sensory integration to facilitate exploration and eventual habituation. The striatum plays a central role in these behaviours, with movement initiation and exploration correlated with an increased in GCaMP fluorescence in both D1- and D2-SPNs^29,44^. Our findings align with these studies, as we observed increased GCaMP7s fluorescence in both D1- and D2-SPNs in WT mice during locomotor responses to a novel environment (Fig. 1c, e). Recent work has further shown that while both D1- and D2- SPNs are active, they participate in distinct neuronal ensembles to select and suppress behaviours via D1-SPN activation and D2-SPN silencing^32,64^. As we measured Ca²⁺ activity across a large dorsal striatal population of direct and indirect SPNs, subpopulation-specific dynamics may be undetected, limiting the degree to which our data align with ensemble-specific models.

Interestingly, in the eIF4E-TG mice, we observed a significant increase in Ca²⁺ activity in D1- SPNs during exploration (Fig. 1c-d), correlating with a heightened novelty response and suggesting cell-type-specific mechanisms underlying hyperactivity and other ASD-related behaviours in the eIF4E-TG mice. Whole-cell recordings further revealed hyperexcitability in D1-SPNs of the eIF4E-TG mice, characterised by a reduction in rheobase (Fig. 2f) and an elevated firing frequency (Fig. 2g-i). These changes indicate that less current is required to initiate action potentials, and once activated, D1-SPNs sustain higher firing rates. This hyperexcitability likely drives the increased Ca²⁺ activity observed *in vivo* during exploration, as reduced rheobase and increased membrane resistance enhance the neuronal responsiveness to novel stimuli. Like FXS models, where altered D1-SPN membrane properties^46^ are linked to locomotor dysregulation^15^, the hyperactive responses of the eIF4E- TG mice likely reflect dysregulated striatal circuits driven by D1-SPN hyperexcitability. In contrast, Ca²⁺ activity in A2a-expressing indirect SPNs during novelty exploration was comparable between eIF4E-TG and WT littermates (Fig. 1e-f). Consistently, no significant changes were detected in excitability parameters of D2-SPNs (Supplemental Fig. 3), though a reduction in capacitance was noted (Supplemental Fig. 3a). Overall, these results demonstrate the cell-type specificity of eIF4E-related alterations and underscore D1-SPN hyperexcitability as a key driver of hyperactive behavioural phenotypes.

To investigate the mechanisms underlying D1-SPN hyperexcitability, we focused on ion channel function, as increased neuronal excitability was associated with changes in membrane resistance, but not in capacitance or neuronal morphology. Specifically, we examined the function of Kir2 and KCNQ potassium channels, both of which are critical regulators of neuronal excitability^65^ and have been implicated in neurodevelopmental disorders, including ASD^66^. We found no dysregulation in Kir2 channel activity in D1-SPNs, indicated by unaltered resting membrane potential (Fig. 2b), excitability during postnatal development (Fig. 3f-i), and Kir2-sensitive currents (Fig. 5a-b)^52^. However, voltage-clamp recordings of D1-SPNs from eIF4E-TG mice revealed a significant reduction in XE991- sensitive currents (Fig. 5c-d), suggesting impaired KCNQ channel function. KCNQ channels, which mediate the M-current, a non-inactivating potassium current activated by depolarisation, are critical for regulating neuronal excitability^65^. Mutations in *KCNQ2*, such as frameshift insertions^67^ and deletions^68^, have been identified in ASD patients. Moreover, conditional deletion of *Kcnq2* in mouse cortical neurons reduces potassium currents and induces hyperexcitability^69^, while heterozygous deletion leads to ASD-like behaviours such as altered exploratory and repetitive behaviours^70^. Conversely, activation of KCNQ channels reduces spontaneous locomotor activity and attenuates dopamine-driven hyperactivity caused by methylphenidate or cocaine^71^. Importantly, our study extends these insights by demonstrating that the reduction in XE991-sensitive currents and the impaired KCNQ activity are cell-autonomous in D1-SPNs of the eIF4E-TG mice. Reducing eIF4E overexpression specifically in adulthood is sufficient to restore KCNQ function and rescue excitability deficits, suggesting a direct link between translational dysregulation and ion channel activity. These results support the hypothesis that translational dysregulation is a molecular mechanism driving ASD pathophysiology^35,72–74^, reinforcing its critical role in shaping neuronal excitability and behavioural outcomes.

The molecular mechanisms underlying the cell-autonomous dysregulation of KCNQ function remain to be determined. KCNQ2 and KCNQ3 subunits can form both homo- and heterodimers, which exhibit distinct biophysical properties, with the latter conveying a larger M-current^75^. If eIF4E affects KCNQ translation in a subunit-specific manner, overexpressing eIF4E could dysregulate translation and result in inappropriate KCNQ subunit composition and function.

Alternatively, the dysfunction in KNCQ channels may arise as part of a mechanism involving D1 receptor signalling. KCNQ channels are closed in response to ERK-dependent phosphorylation driven by D1 receptor activation, which increases D1-SPN excitability^60^. Our recent findings demonstrated that eIF4E-TG mice exhibit reduced dopamine release in the striatum^41^, which could potentially result in a compensatory increase in D1 receptor expression. In addition, eIF4E itself is phosphorylated via ERK signalling through Mnk1/2 kinases, enhancing cap-dependent translation^76,77^. Altogether, these findings raise the intriguing possibility that increased D1 receptor signalling leads to enhanced ERK-mediated phosphorylation of both KCNQ channels and eIF4E, which could synergistically amplify the effects of aberrant translation and contribute to the hyperexcitability of D1-SPNs.

Another possible mechanism involves dysregulated passive membrane properties in D1- SPNs from early developmental stages, leading to compensatory changes in evoked firing properties later in life. During early postnatal development, SPNs do not exhibit their characteristic low resting membrane potential (RMP) due to minimal Kir2 channel activity, which increases during maturation^52^. However, the D1-SPNs of eIF4E-TG mice exhibit a hyperpolarised RMP already at P10 (Fig. 3b). This suggests that, unlike evoked firing properties, passive membrane properties are disrupted from birth. This early hyperpolarised state may trigger homeostatic changes in K+ channel dynamics, such as altered KCNQ function, ultimately resulting in increased firing frequency.

Our results emphasise that the role of KCNQ channels in striatal development should be further explored, particularly in the context of therapeutic development for neurodevelopmental disorders characterised by neuronal hyperexcitability. Drugs that promote KCNQ channel function, such as retigabine (ezogabine), reduce epilepsy and are in clinical trials for various neurological disorders^78–80^. It will be essential to evaluate whether such compounds are effective in ameliorating the hyperexcitability phenotypes and associated behaviours in our eIF4E-TG model.

An alternative therapeutic option could be targeting translational regulation pathways, particularly the eIF4E signalling axis. Several studies highlight the potential for postnatal mTOR inhibition in reducing ASD pathology. For instance, in VPA-treated rodents, postnatal administration of rapamycin, an mTOR inhibitor, successfully reduced SPN hyperexcitability and aberrant firing patterns^81^. Furthermore, studies in the Tsc1 and Tsc2 models of tuberous sclerosis have demonstrated the efficacy of postnatal mTOR inhibition in ameliorating diverse ASD-like phenotypes^82^, ranging from cognitive and social impairments in heterozygote *Tsc2^+/-^* mice^83,84^ to morphological abnormalities and survival deficits in *Tsc1* deletion models^85^, and epilepsy in mice with glia-specific *Tsc1* deletion^86^. The fact that D1-SPN hyperexcitability (Fig. 2f-i) in eIF4E-TG mice emerges after the main postnatal developmental period (Fig. 3f-i) and, along with hyperactivity (Fig. 6a-b) and repetitive behaviours (Fig. 6c) is normalised with eIF4E reduction, suggests that interventions targeting eIF4E- and mTOR-related pathways to treat ASD-like motor behaviours may remain effective in adulthood.

Given the cell-type-specific nature of the observed hyperexcitability, interventions selectively targeting D1-MSNs or the eIF4E pathway may offer significant advantages over more generalised approaches by minimising off-target effects. Our successful application of conditional RNAi to reduce eIF4E expression provides proof-of-concept for the development of RNAi-based therapies or small molecules designed to modulate eIF4E activity or its downstream signalling pathways^87^. Interestingly, we observed no severe effects of eIF4E reduction in WT mice, highlighting a potentially favourable safety profile for such targeted interventions. These findings suggest that reducing eIF4E expression can effectively rescue pathological phenotypes in eIF4E-TG mice while preserving normal neuronal and behavioural function in WT mice, emphasising the therapeutic potential of postnatal interventions targeting eIF4E signalling.

In conclusion, our findings broaden the understanding of translational dysregulation in ASD, demonstrating that its impact extends beyond previous findings of synaptic dysfunction^40–43^ to encompass intrinsic excitability changes in striatal neurons. By showing that translational dysregulation in D1-SPNs affects excitability and potassium channel function, our study provides compelling evidence that mRNA translation is a central regulator of neuronal activity. Furthermore, genetic reduction of eIF4E in adult D1-SPNs ameliorates excitability, potassium channel function, and notably hyperactivity and repetitive behaviours. These results highlight the potential of modulating translational pathways as a targeted, cell-specific therapeutic strategy to address the complex neuronal phenotypes associated with ASD.

## Materials and methods

### Mice

All animal experiments were compliant with the ethical permit issued by the Swedish Board of Agriculture (Ethical number: 18194-2018 and 19345-2023) and were performed in accordance with the European Parliament and Council Directive 210&63/EU, 22nd September 2010 for experimentation animals’ protection. Mice were sex-separated and group-housed (up to five mice per cage) in a temperature (23C) and humidity (55%) controlled environment, on a 12-h light/dark cycle with water and food available *ad libitum*. In all cases, eIF4E^wt/wt^ mice are referred to as wild type (WT), and eIF4E^wt/**β**tEif4e^ mice are referred to as eIF4E-transgenic (eIF4E-TG or simply TG)^40,88^. Each of the following lines were maintained in hemizygosis by crossing with C57BL/6J mice purchased from Janvier and were crossed with the eIF4E-TG line to generate double transgenic mice. All mice were maintained on the C57BL/6J genetic background.

For the fibre photometry experiment, we used three-to-five-month-old male double-transgenic mice generated by crossing Tg(Drd1a-cre)FK150Gsat^+/-^ (D1-Cre) or Tg(Adora2a- cre)KG139Gsat^+/-^ (A2a-Cre) mice^89^ with eIF4E^wt/**β**tEif4e^ , producing D1-Cre/WT, D1-Cre/TG, A2a-Cre/WT and A2a-Cre/TG to target D1- and D2-SPNs, respectively. For intrinsic and K+ channel recordings, we used postnatal day 60-90 male double-transgenic mice generated by crossing Drd1a-TdTomato ^+/-^ (D1-Tomato) mice^47^ (Jackson Laboratory, #016204) with eIF4E^wt/**β**tEif4e^ producing D1-Tomato/WT and D1-Tomato/TG. For developmental recordings, we used postnatal day 10 (P10) and 28 (P28) mice, both males and females, of the same strain. For the neuronal morphology experiments, we used adult double-transgenic mice generated by crossing Tg(Drd2-EGFP)S118Gsat^+/-^ (D2-EGFP) mice ^90^ with eIF4E^wt/**β**tEif4e^ yielding D2- EGFP/WT and D2-EGFP/TG. For eIF4E reduction experiments (RNAi), we used two-to-four-month-old male double-transgenic mice generated by crossing TRE-GFP.shmiR-4E mice^55,91^ with eIF4E^wt/**β**tEif4e^, generating TRE-GFP.shmiR-4E/WT and TRE-GFP.shmiR-4E/TG mice.

### Fibre photometry

Fibre photometry measurements were conducted using custom-built optical components and custom LabVIEW software^92^. The setup included two fibre-coupled LEDs: one for exciting Ca²⁺-bound GCaMP7s (M470F3, Thorlabs) modulated at 573Hz, and the other for exciting the isosbestic signal (M405F1, Thorlabs) modulated at 211Hz. The LED beams passed through emission filters (FF02-472/30-25, Semrock; FB405-10, Thorlabs) and a long pass dichroic mirror (DMLP425R, Thorlabs). They were then reflected via a dichroic beamsplitter (FF495- Di03-25x36, Semrock) and directed through a fibre-optic patch cord (400µm core, 0.5NA, R- FC-P-N5-400-L1, RWD Life Science) connected to the mouse using a ceramic mating sleeve (ADAF1-5, Thorlabs). Fluorescence emitted from the sample was collected through the same patch cord, filtered by a bandpass filter (FF01-535/50-25, Semrock), and focused onto the sensor of a battery-powered femtowatt photoreceiver (Newton, Model 2151) using a plano-convex lens (62-561, Edmund Optics). The excitation LED modulation and lock-in amplification of the photoreceiver signal were managed by a Field-Programmable Gate Array board (National Instruments, sbRIO-9637). The light power was set to 50µW for both LEDs.

Mice were habituated to a cage (Tecniplast GM500, 500cm^2^) across two days prior to recording. On the day of recording, mice were introduced to the same cage for at least 30 minutes, after which the baseline GCaMP7s fluorescence was collected. After five minutes, the mice were transferred to a novel cage environment and the GCaMP7s activity was monitored for 10 minutes. In a separate experiment, mice were intraperitoneally (i.p.) injected with D1-receptor blocker SCH39166 (0.25mg/kg, Tocris) during recording to assess the effect of D1-receptor blockade on GCaMP7s activity.

### Analysis

Recordings were analysed using a custom-made MATLAB script. For the pre-processing we used steps similar to those previously described^93^. In brief, we applied a low-pass Butterworth filter of 25Hz and performed exponential curve fitting to both the GCaMP data signal (470nm) to the isosbestic (405nm) control. We then fitted the GCaMP signal to the isosbestic control using scaled linear regression as described^93–95^. We performed Z-score normalisation of the fitted signal prior to analysis.

We then quantified signal peak frequency and size from one-minute recording sections of baseline and novelty-induced responses, to detect small scale changes in GCaMP signal – similar to parameters measured previously during longitudinal fibre photometry recordings^96,97^. For these, we used the Find Peaks function in MATLAB to quantify the frequency of peaks in Hz in the given time frame, and we used the envelope function to measure the amplitude of the signal by quantifying the difference between the average upper and lower limits of the Z-score fluctuations during the same time frame. We also analysed the area under the curve of the one-minute recordings of the signal to detect bulk changes in GCaMP activity. These three parameters (frequency, size and area under curve) were normalised to their respective baseline values and then combined to create an aggregate ‘composite’ score, which provides an overall, multi-parameter change in GCaMP7s activity, expressed as a percentage from baseline (0%).

### Stereotaxic surgeries

Mice were injected with buprenorphine (i.p. 0.1mg/kg) and anaesthetised with 2% isoflurane. Adenovirus (AAV) injection was administered via a Quintessential Stereotaxic Injector (Stoelting) with a 30nl/min flow rate. Post-operative treatment was given in the form of carprofen and buprenorphine i.p. injection (5mg/kg and 0.1mg/kg, respectively). Mice were left to recover for at least three weeks prior to experimentation, to allow sufficient viral expression.

For the fibre photometry experiment, unilateral injection of 250nl pGP-AAV-*syn*-FLEX- jGCaMP7s-WPRE (Addgene #104491) and implant of fibre optic cannula (RWD Life Sciences: 0.5NA, 400µm core, 2.5mm) were performed in the dorsal striatum (AP +0.8, ML -1.8, DV -2.4 taken from dura). In separate control experiments, mice were injected with 250nl pAAV-*hSyn*- EGFP (Addgene #50465) to confirm the absence of non-GCaMP mediated events.

Selective eIF4E reduction (RNAi) was performed similarly to previously described methods^53,91^. For the initial characterisation of eIF4E reduction in SPNs, we performed bilateral injections of 300nl AAV5*-hSyn-*mCherry*-*Cre (UNC Vector Core) and 300nl pAAV-*ihSyn1*-DIO-tTA (Addgene #99121) or 300nl pAAV-FLEX-tdTomato (Addgene #28306) in the dorsal striatum. To specifically target D1-SPNs, bilateral injections of 150nl pAAV-*EF1a*-*Cre* (Addgene #55636) in the substantia nigra pars reticulata (SNr) (AP -3.28, ML +/-1.5, DV -4.3) were performed to retrogradely target direct pathway neurons, i.e., D1-SPNs, with bilateral striatal injection of 300nl pAAV-*ihSyn1*-DIO-tTA (Addgene #99121) or 300nl pAAV-FLEX-tdTomato (Addgene #28306). For the intrinsic property recordings, we bilaterally injected 150nl pAAV-*EF1a*-*Cre* (Addgene #55636) in the SNr and unilaterally injected 300nl pAAV-*ihSyn1*- DIO-*tTA* (Addgene #99121) in the left dorsal striatum (AP +1 and 0, ML +1.8, DL -2.4) and 300nl pAAV-FLEX-tdTomato (Addgene #28306) in the right striatum (AP +1 and 0, ML -1.8, DL -2.4). For the KCNQ and behavioural experiments, we bilaterally injected either the pAAV-FLEX-tdTomato (Addgene #28306) or pAAV-*ihSyn1*-DIO-*tTA* (Addgene #99121) in the dorsal striatum.

### Electrophysiology

#### Slice preparation

Acute brain slices were prepared and maintained using the two-step *N-*methyl-D-glucamine (NMDG) based protective recovery method^98^. Slices were initially prepared using NMDG-based solution containing (in mM): 92 NMDG, 30 NaHCO3, 2.5 KCl, 20 HEPES, 2 thiourea, 1.25 NaH2PO4, 3 Na-pyruvate, 5 ascorbic acid, 25 glucose, 0.5 CaCl2 and 10 MgCl2 titrated to pH 7.4. During the experiment, the slices were stored in the HEPES-based holding solution containing (in mM): 92 NaCl, 30 NaHCO3, 2.5 KCl, 20 HEPES, 2 thiourea, 1.25 NaH2PO4, 3 Na-pyruvate, 5 ascorbic acid, 25 glucose, 2.5 CaCl2 and 2 MgCl2 titrated to pH 7.4. Briefly, mice underwent cardiac perfusion with 25ml chilled NMDG solution prior to brain removal and dissection. 250µm coronal slices were collected containing the dorsal striatum and stored in NMDG-based solution for 10 minutes at +32C before transferring to HEPES-based holding solution. All solutions were oxygenated with 95% O2 and 5% CO2.

#### Recording protocols

All recordings were obtained using a potassium-gluconate based internal solution containing (in mM): 120 K-gluconate, 20 KCl, 4 MgATP, 0.3 Na2-GTP, 5 Na2-phosphocreatine, 0.1 EGTA and 10 HEPES at pH 7.25 and osmolarity ∼300mOsm, with a pipette resistance of 3-5MΩ. During the experiments, slices were placed in the submersion recording chamber in the rig with oxygenated (95% O2/ 5% CO2) artificial cerebrospinal fluid (ACSF) supplied at 2-3ml/min and maintained at +32-34C, containing (in mM): 125 NaCl, 2.5 KCl, 25 NaHCO3, 1.25 NaH2PO4, 10 glucose, 2 CaCl2 and 1 MgCl2. Potentials were not corrected for the liquid junction potential. For recordings of Kir2 channel activity, ACSF was prepared with 300nM tetrodotoxin citrate (TTX, Tocris) to prevent depolarisation and 1mM CsCl2 to block potassium channels. For recordings of Kv7 channel activity, we used 10µM XE991-dihydrochloride (Tocris) in ACSF.

D1- and D2-SPNs were identified based on presence or absence, respectively, of TdTomato fluorescent marker. Intrinsic properties were recorded in current-clamp configuration. Ramp current injection was performed ranging from -300 to +450pA over one second with a one-second inter-sweep interval. The final recording was performed as stepwise current injection from -300pA to +450pA in 50pA increments. Each current step was held for 500ms with 500ms inter-sweep interval. The Kir2 channel recordings were performed in voltage-clamp configuration, as previously described^52^. Cells were held at -60mV and voltage steps were performed in -10mV increments until -140mV, after which CsCl2 (1 mM) was bath-applied, and the voltage steps were performed again 10 minutes after bath application. To activate the M-current, we clamped the membrane potential at 0mV for four seconds, followed by a series of repolarising, deactivating voltage steps from -20mV to -60mV, before and after application of XE991 to block KCNQ2/3 channels^60^. For all recordings, the access resistance was monitored throughout the recording by applying hyperpolarizing 10mV voltage steps. Recordings were discarded if access resistance exceeded 30MΩ or changed by over 10%.

#### Analysis

All analysis of electrophysiological data was performed using the open-access Python-based software Stimfit^99^. The current at which the first action potential was fired was used to determine the rheobase current. The rheobase action potential was used to calculate the action potential height and half-height duration. Current-frequency (IF) plots were generated from the action potential frequency (Hz) at each current step. Membrane capacitance was calculated using 𝞃/Ri, where Ri is the input resistance calculated from the voltage response to -100pA current injection, and 𝞃 is the membrane time constant, calculated using an exponential fit from the initial voltage change with -100pA current. The membrane resistance (Rm) was determined using Ohm’s law (V=IR) for the range of current steps performed. For Kir2 and KCNQ channel recordings, the drug-sensitive current was calculated as the difference between the current responses before and after CsCl2 or XE991 application.

### Behaviour

#### Novelty induced locomotor activity

Mice were individually placed in clean homecages (Tecniplast GM500, 501cm^2^) with minimal bedding material covering the bottom and allowed to explore for two hours. The novelty-induced locomotor activity test was conducted in a dark environment under infrared lighting between 9AM and 5PM. During the test, mice were video recorded, and their distance travelled was analysed using Ethovision XT16 video tracking software (Noldus, The Netherlands).

#### Marble Burying

For the marble burying test, 20 marbles were evenly arranged in a large cage (Tecniplast GR900, 904cm^2^) with at least 5cm of fine wood shavings. Mice were introduced to the cage and allowed to move freely for 30 minutes. After this period, the number of marbles buried was scored, where a marble was considered buried if less than half of it was visible above the surface.

#### Western blotting

Mice were sacrificed via cervical dislocation and striata from the left and right hemispheres were dissected and flash-frozen in liquid nitrogen. The samples were sonicated in 1% SDS and boiled for 10 minutes^50^. Protein concentration was determined using the Pierce™ BCA Protein Assay Kit. A standard curve was generated from a colorimetric assay of bovine serum albumin (BSA) with known concentrations to quantify the protein levels in our samples. Samples were diluted in Laemmli buffer (4X) and boiled for one minute. Equal quantities of protein were loaded (5µg or 30µg) into 10% polyacrylamide gels. Proteins were transferred to Immobilon FL PVDF membranes (pore size 0.2µm). Membranes were blocked for one hour in 5% milk in TBS with 0.5% Tween-20 (TBS-T) before incubation with primary antibody for two hours at room temperature (R&D system MAB3228 mouse anti-eIF4E, Cell Signalling 2306S rabbit anti-DARPP32; diluted 1:20,000 and 1:50,000, respectively). Blots were washed in TBS- T and incubated in secondary antibody (Invitrogen A16096 anti-rabbit or A16066 anti-mouse HRP conjugated antibody, diluted 1:50,000 and 1:25,000, respectively) for 30 minutes. Following further washes in TBS-T, blots were treated with Pierce ECL Plus Western Blotting Substrate for five minutes and developed in a dark room using a Protec OPTIMAX X-Ray Film Processor. The resulting films were scanned, and the images were analysed using ImageJ to quantify the protein expression levels.

#### Immunofluorescence

Mice were sacrificed and transcardial perfusion with 4% paraformaldehyde was performed. Whole brains were extracted and 30-40µm coronal slices were prepared with a vibratome in PBS. Slices containing striatum were selected for immunostaining. Slices were washed in TBS before permeabilization for five minutes in solution containing 1:10 methanol and 1:3 hydrogen peroxide in TBS. A second permeabilization step was performed using TBS with 0.2% Triton X-100 (TBS-Triton) and incubated for 15 minutes. Slices were washed in TBS and then incubated in primary antibody overnight at +4C (1:200 R&D system MAB3228 mouse anti- eIF4E, 1:500 Synaptic System 382004 guinea pig anti-DARPP32, 1:400 ImmunoStar 20065 rabbit anti-enkephalin, 1:2000 Abcam ab13970 chicken anti-GFP, 1:1000 Rockland 600-401- 379 rabbit anti-RFP, 1:500 Rockland 200-301-379 mouse anti-RFP). Slices were washed and then incubated in secondary antibody for 45-60 minutes at room temperature (Invitrogen Alexa fluor: A-11001 anti-mouse 488, A-11004 anti-mouse 548, A-21450 anti-guinea pig 647, A-11039 anti-chicken 488, A-11008 anti-rabbit 488, A-11011 anti-rabbit 568; all diluted 1:500). Slices were mounted onto glass slides with Fluromount-G with DAPI (Invitrogen). Sections were imaged at 10x or 20x with a Zeiss LSM800 confocal microscope, and images were processed with ImageJ for manual counting.

#### Neuronal morphology

The process of microinjection was followed as previously described^100^. Briefly, mice underwent cardiac perfusion, and brains were removed and fixed for four hours in 4% paraformaldehyde (PFA) in 0.1M PBS (pH 7.4) at 4°C. Brains were sliced at 200µm and stored in PBS with 0.1% Na-azide. During microinjection, D2-SPNs were identified via expression of GFP and D1- SPNs via absence of GFP, and cells were filled with 8% Lucifer Yellow CH lithium salt (Invitrogen) in PBS and Tris-HCl via current injection of 2-10nA over five-10 minutes. Cells were filled until distal dendrites were visible, and post-fixed in PFA before imaging. To amplify the fluorescent signal from Lucifer Yellow, following a series of washes in tris-buffered saline (TBS), slices were incubated in Streptavidin Alexa Fluor™-488 (Invitrogen, 1:200) in 0.6% Triton X-100 in TBS for 48 hours. Slices were then incubated overnight in 2% NGS / 0.6% TBS-triton with goat anti-mouse Alexa Fluor™ 488 (Invitrogen, 1:500). Cell morphology was analysed using the Sholl analysis function of the Neuroanatomy plug-in for ImageJ.

### Statistical analysis

All statistical analyses were performed using Graphpad Prism. For behavioural, fibre photometry and western blot experiments, the n-values indicate the number of mice, and data are presented as mean ± SEM. For electrophysiology and neuronal morphology experiments, the N-values indicate number of mice and n-values indicate the number of cells and data are presented as median ± quartiles, whereas for immunofluorescence experiments the n-values indicate the number of slices. Either two-tailed student t-tests or paired t-tests were used for two-group comparisons, whereas two-way ANOVAs were used for grouped variable comparisons and repeated-measures (RM) two-way ANOVA for matched variable comparisons, with Tukey’s multiple comparisons test for post-hoc comparisons between groups. When comparing three parameters, a three-way ANOVA was used. Multiple comparisons tests were performed only in the case of an interaction effect, and comparisons were made only between genotypes in the same condition, or between conditions in the same genotype. In all cases, * p<0.05, ** p<0.01, *** p<0.001 and **** p<0.0001, ns p>0.05. Key statistical parameters, including N and n values, mean ± SEM, median ± quartiles, t- and f-values, degrees of freedom, p-values and significance levels are indicated for each statistical test in supplemental tables 1-12.

## Acknowledgements

We thank the members of Borgkvist and Santini labs for their invaluable methodological support and insightful discussions. We extend special thanks to Eric Klann for generously providing the founders of the eIF4E^wt/**β**tEif4e^ and TRE-GFP.shmiR-4E mice and Robert Fetcho for designing the fibre photometry system. We are also grateful to Qian Yu and the KI Animal Behavioral Core Facility for their support during the behavioural experiments, as well as to the staff and veterinarians of the Comparative Medicine Biomedicum (KM-B) for their continuous assistance in maintaining the mouse colonies. This work was supported by the Knut and Alice Wallenberg Foundation (Wallenberg Academy Fellow Grant KAW 2017-0169 and project grant 2020-0054 to E.S.), the Swedish Research Council (2016-02758 and 2023-02943 to E.S. and 2016-03129 to A.B.), Olle Engkvist Stiftelse (E.S. and A.B.), Åhlén’s foundation (A.B.), Magn. Bergvall’s foundation (A.B.), The Strategic Research Program in Neuroscience (StratNeuro) starting (E.S. and A.B.) and bridging (E.S.) grants, Karolinska Institute starting grant (E.S.).

## Author contributions

Conceptualisation, A.B. and E.S.; Methodology, A.A., A.B. and E.S.; Validation, A.A., A.B. and E.S.; Formal Analysis, A.A., A.B. and E.S.; Investigation, A.A., A.T., A.P.R., A.B. and E.S.; Resources, A.B. and E.S.; Writing – Original Draft, A.A., A.B. and E.S.; Writing – Review & Editing, A.A., A.B. and E.S.; Visualisation, A.A, A.B. and E.S.; Supervision, A.B. and E.S.; Project Administration, A.B. and E.S.; Funding Acquisition, A.B. and E.S.

