## Supplementary figures and tables for "Postnatal reduction of eIF4E overexpression in D1-SPNs ameliorates KCNQ dysfunction, hyperexcitability and ASD-like behaviours"

### Supplemental Information

#### Supplemental figures

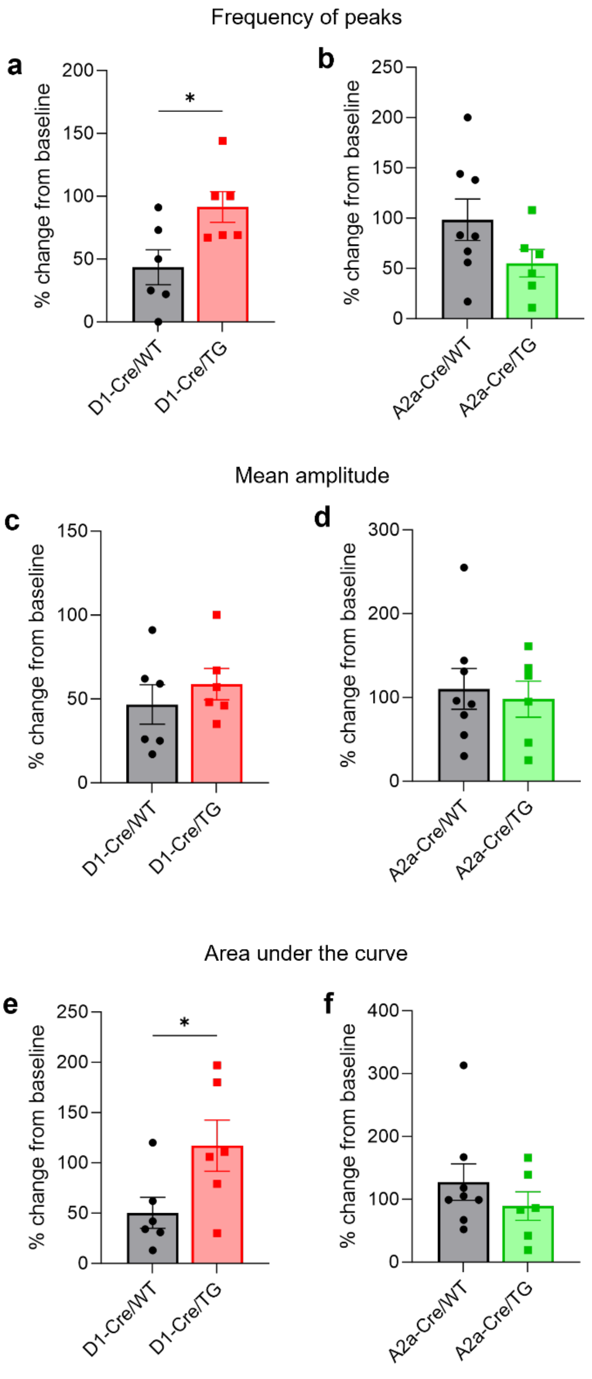


#### Supplemental figure 1: GCaMP7s analysis parameters in the direct and indirect pathway SPNs during a novelty conditioning paradigm.

**a-b**) change in peak frequency in the D1-Cre (unpaired T-test: t=2.581, *p=0.0274) and A2a-Cre mice. **c-d**) change in mean signal amplitude in the D1-Cre and A2a-Cre mice. **e-f**) change in area under the curve (AUC) in the D1-Cre (unpaired T-test: t=2.243, *p=0.0487) and A2a-Cre mice. In all panels, normalised changes in

GCaMP7s signals were obtained from 6 D1-Cre/WT and 6 D1-Cre/TG and 8 A2a-Cre/WT and 6 A2a-Cre/TG mice. Values are expressed as percentage changes from baseline. Data are presented as mean ± SEM, with mean indicated by bar height and SEM by error bars. Dots represent individual values for each mouse. For full details of statistical analysis including negative results, refer to supplemental table 8.

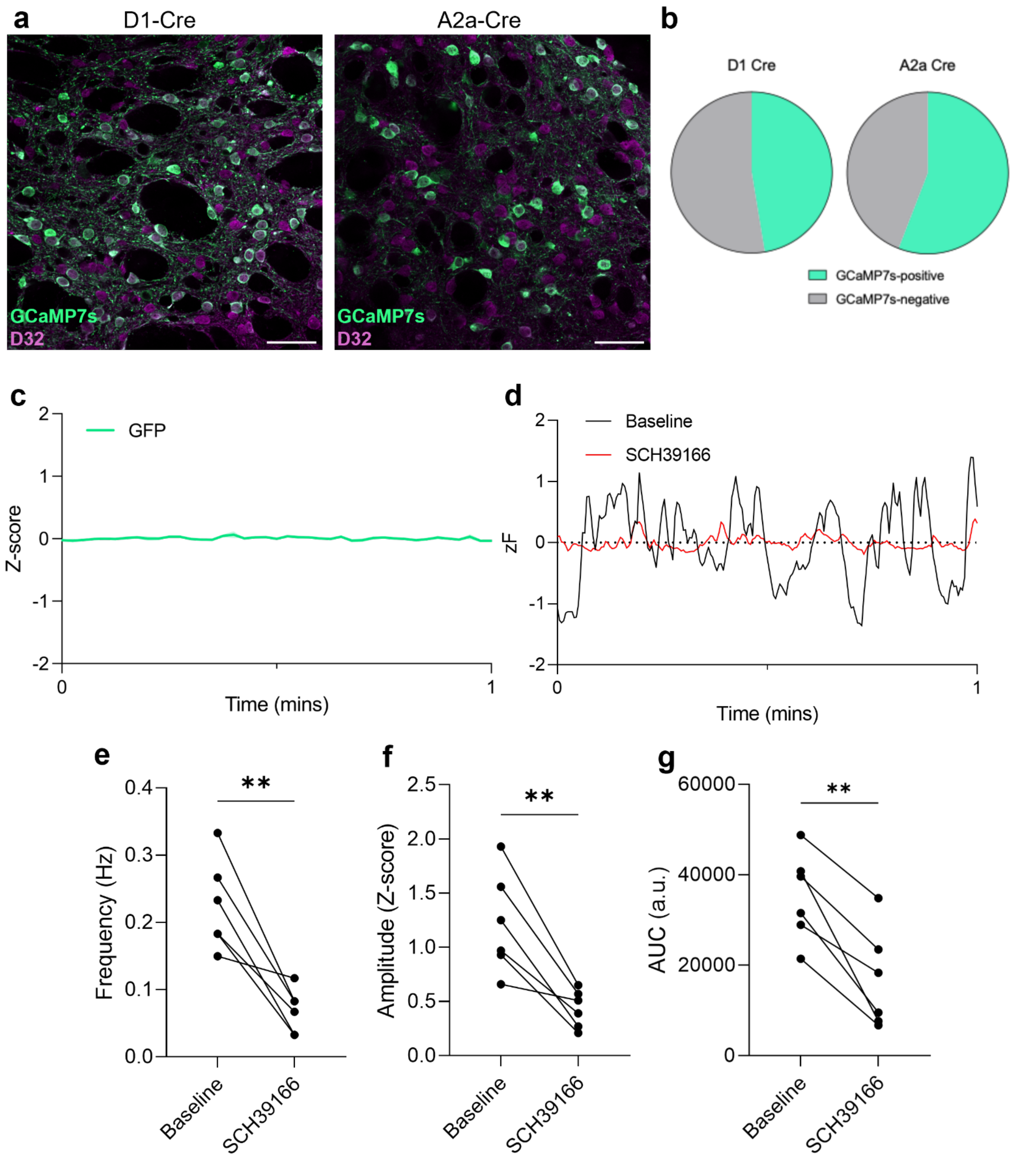

#### Supplemental figure 2: Pathway specificity of GCaMP7s activity.

**a**) Representative confocal images showing the striatal distribution of GCaMP7s-expressing cells (green) in D1-Cre (left) and A2a-Cre mice (right), with DARPP32 immunostaining in magenta (scale bar: 50µm). **b**) Proportion of GCaMP7s-positive cells in striatal brain slices from D1-Cre and A2a-Cre mice. **c**) Representative trace of fibre photometry recording from striatal GFP-injected mice over a one-minute period. **d-g**) Effect of the D1-receptor antagonist SCH39166 on GCaMP7s activity. **d**) Representative traces showing GCaMP7s activity at baseline (black) and 15 minutes after intraperitoneal (i.p.) injection of SCH39166 (DOSE; red). Both traces were quantified using the same Z-score normalisation. To allow superimposition, the mean of each one-minute trace further normalised to zero. **e**-**g**) SCH39166 (DOSE, i.p.) significantly reduces GCaMP7s signal 15 minutes after injection affecting all three parameters : **e**) peak frequency in hertz (paired T-test: t=5.062, **p=0.0039), **f**) signal amplitude, expressed as a z-scored values (paired T-test: t=4.875, **p=0.0046), and **g**) area under the curve (AUC) of the signal in arbitrary units (a.u.; paired T-test: 5.554, **p=0.0026). Each data point represents individual recordings before and after drug administration (6 D1-Cre/WT mice). For panels **e-**g), data is shown as raw values without normalisation to baseline. For full details of statistical analysis including negative results, refer to supplemental table 7.

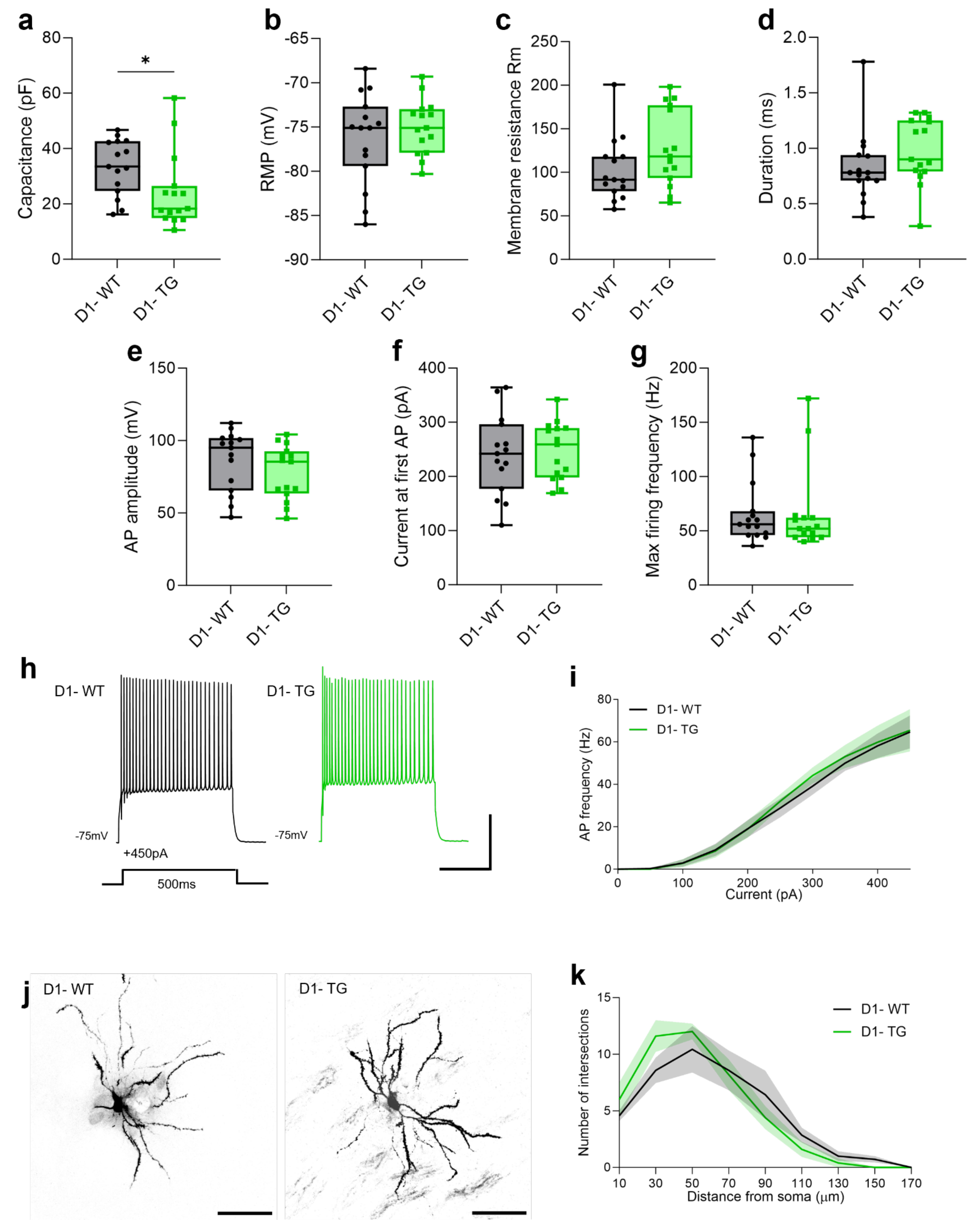


#### Supplemental figure 3: Ex vivo recordings of intrinsic properties in D2-SPNs.

**a**) Calculated capacitance in D2-SPNs (D1- cells) of D1-TdTomato/WT and -/TG mice (unpaired T-test: t=2.062, *p=0.0486). **b**) Resting membrane potential. **c**) Membrane resistance. **d**) Action potential half-height duration. **e**) Action potential peak amplitude. **f**) Rheobase current. **g**) Maximum firing frequency across the range of injected currents. **h**) Representative traces of action potentials at +450pA current injection for 500ms (scale bar: 200ms, 40mV). **i**) Current-frequency (IF) plots for D2-SPNs. **j**) Representative images of Lucifer Yellow-filled D2-SPNs (scale bar: 50μm). **k**) Dendritic intersections plotted by distance from soma. Data are presented as median ± quartiles within the box plot, and whiskers indicate minimum and maximum values (**a**-**g**), or as mean ± SEM, with mean indicated by the connecting line and SEM by error lines with shading (**i** and **k**). Dots represent individual values for each mouse. All recordings were collected from D1- cells from 5 D1-TdTomato/WT and 5 D1-TdTomato/TG mice (n=15 WT, n=15 TG). For full details of statistical analysis including negative results, refer to supplemental table 9.

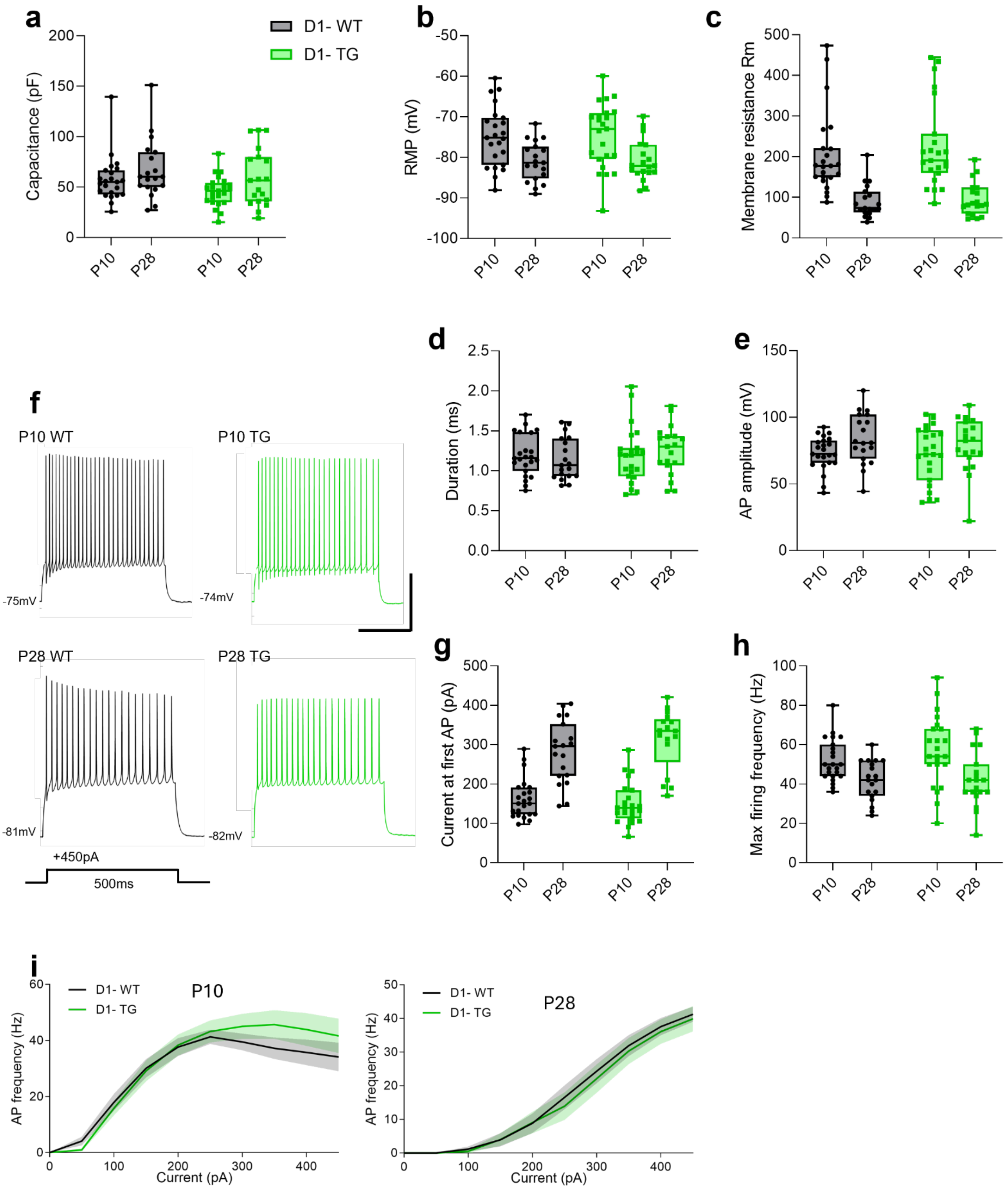


#### Supplemental figure 4: Intrinsic properties of D2-SPNs across postnatal development.

**a**) Calculated capacitance of D2-SPNs from D1-Tomato/WT and D1-Tomato/TG at P10 and P28. **b**) Resting membrane potential of D2-SPNs at P10 and P28 mice. **c**) Membrane resistance of D2-SPNs at P10 and P28 mice. **d**) Action potential half-height duration of D2-SPNs at P10 and P28 mice. **e**) Action potential peak amplitude of D2-SPNs at P10 and P28 mice. **f**) Representative traces of action potentials at +450pA current injection for 500ms (scale bar: 200ms, 50mV). **g**) Rheobase current of D2-SPNs at P10 and P28 mice. **h**) Maximum firing frequency across the range of injected currents of D2-SPNs at P10 and P28 mice. **i**) Current-frequency (IF) plots of D2-SPNs at P10 and P28 mice. Data are presented as median ± quartiles within the box plot, and whiskers indicate minimum and maximum values (**a-h**), or as mean ± SEM, with mean indicated by the connecting line and SEM by error lines with shading (**i**). Dots represent individual values for each mouse. All recordings were collected from D1- cells from 7 D1-TdTomato/WT and 5 D1-TdTomato/TG P10 mice and 5 D1-TdTomato/WT and 5 D1-TdTomato/TG P28 mice (n=23 WT P10, n=23 TG P10, n=19 WT P28, n=19 TG P28). For full details of statistical analysis including negative results, refer to supplemental table 10.

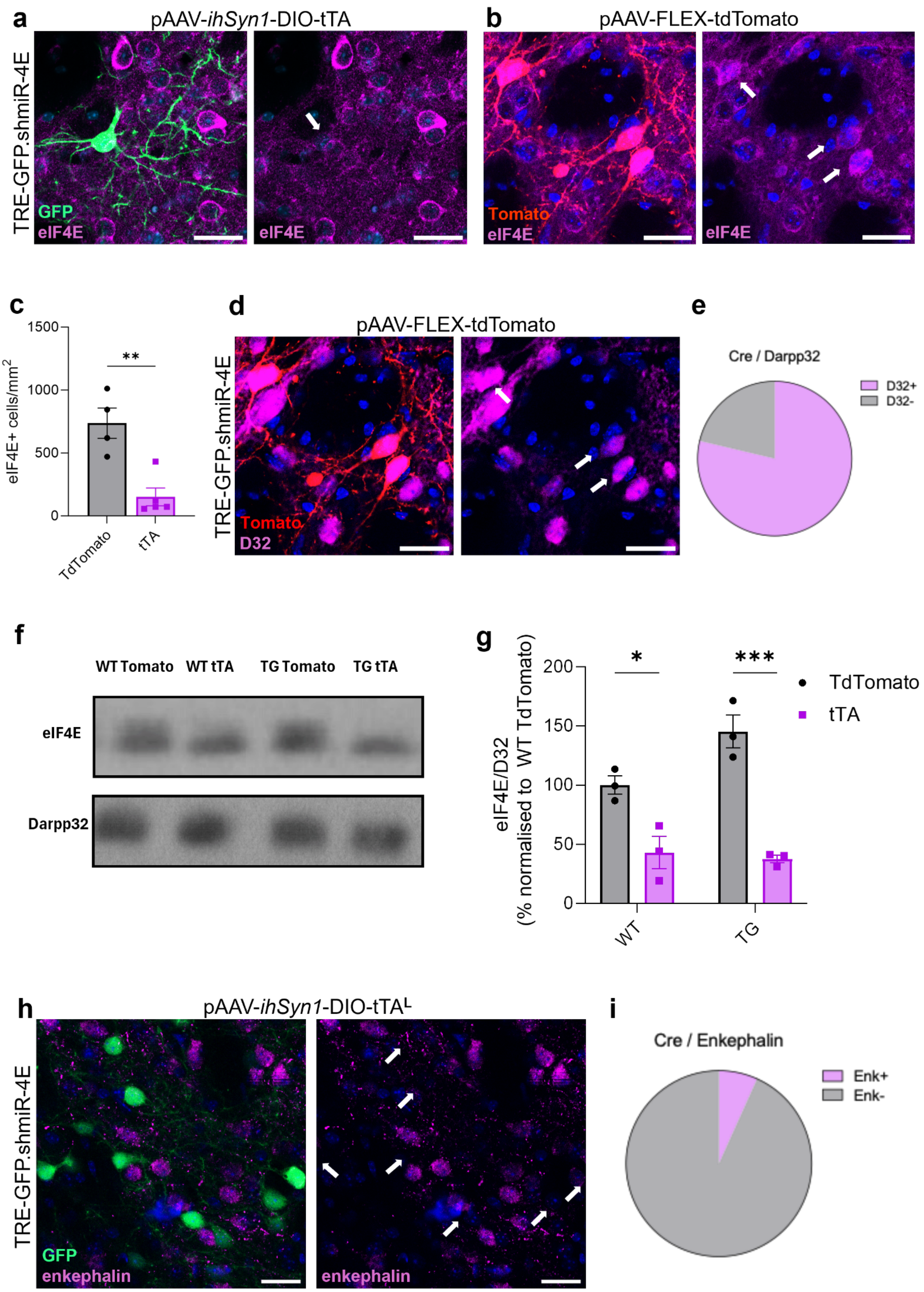
a
pAAV-ihSyn1-DIO-tTA
TRE-GFP.shmiR-4E
GFP
eIF4E
eIF4E
b
pAAV-FLEX-tdTomato
Tomato
eIF4E
eIF4E
c
eIF4E+ cells/mm2
1500
1000
500
0
**
TdTomato
tTA
d
pAAV-FLEX-tdTomato
TRE-GFP.shmiR-4E
Tomato
D32
e
Cre / Darpp32
D32+
D32-
f
WT Tomato
WT tTA
TG Tomato
TG tTA
eIF4E
Darpp32
g
eIF4E/D32
(% normalised to WT TdTomato)
200
150
100
50
0
*
***
TdTomato
tTA
WT
TG
h
pAAV-ihSyn1-DIO-tTAL
TRE-GFP.shmiR-4E
GFP
enkephalin
enkephalin
i
Cre / Enkephalin
Enk+
Enk-

#### Supplemental figure 5: Conditional RNAi system to reduce eIF4E expression.

**a**) Representative confocal image of the striatum showing the reduction of eIF4E immunostaining (magenta) in GFP-expressing neurons (green) of TRE-GFP.shmiR-4E mice striatally injected with AAV-Cre together with an AAV-DIO-tTA. **b**) Representative confocal image showing eIF4E expression (magenta) in TdTomato-expressing neurons (red) of TRE-GFP.shmiR-4E mice striatally injected with AAV-Cre with an AAV-FLEX-TdTomato. **c**) Quantification of the striatal density of eIF4E-positive cells in TdTomato and tTA-injected mice (unpaired t-test: t=4.417, **p=0.0031). **d**) Representative images showing the DARPP-32 expression (magenta) in TdTomato-expressing neurons (red). **e**) Pie chart depicting the proportion of Cre-expressing (either TdTomato- or GFP-positive) cells that colocalise with DARPP-32 immunostaining. **f**) Representative Western blot images for the level of eIF4E in TRE-GFP.shmiR-4E/WT and -/TG TdTomato and tTA-injected striatal extracts, with DARPP-32 as a loading control. **g**) quantification with each sample normalised to their respective loading control and expressed as percentage of the TRE-GFP.shmiR-4E/WT TdTomato (genotypeXRNAi interaction, 2-way ANOVA: F(1, 8)=5.836, *p=0.0421). **h**) Representative images showing enkephalin immunostaining (magenta) compared with GFP expression (green). **i**) Pie chart depicting the proportion of Cre-expressing cells (from either TdTomato$^{R}$ or tTA$^{L}$ slices) that colocalise with enkephalin immunostaining. In all the confocal images, DAPI is shown in blue and scale bars indicate 25µm. Data are presented as mean ± SEM, with mean indicated by bar height and SEM by error bars. Dots represent individual values for each mouse. Significance is denoted as * p<0.05, ** p<0.01, *** p<0.001 and **** p<0.0001, calculated with Tukey's post-hoc multiple comparisons test. For full details of statistical analysis including negative results, refer to supplemental table 11.

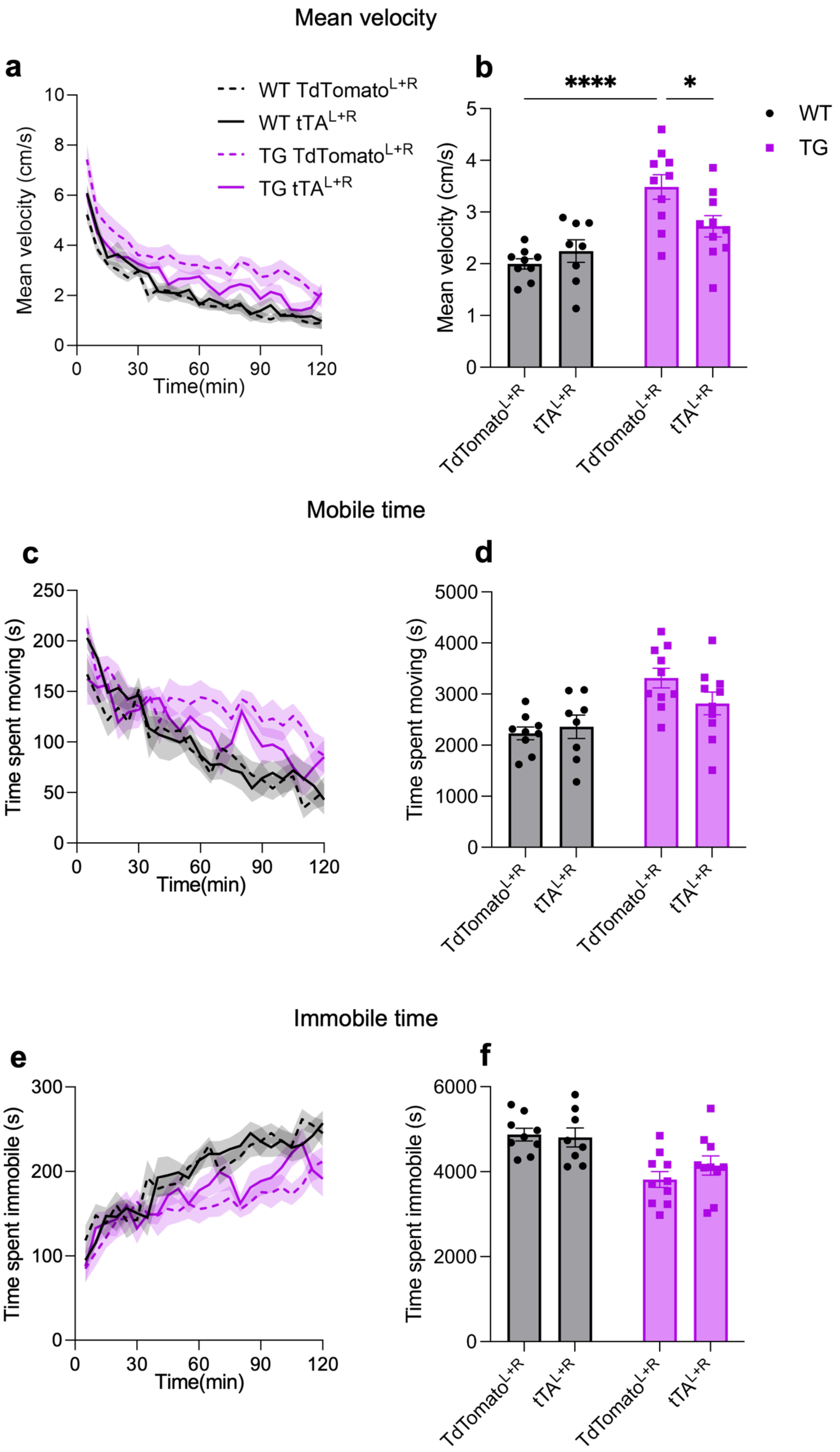
Mean velocity
a
b
WT TdTomato L+R
WT tTA L+R
TG TdTomato L+R
TG tTA L+R
Mean velocity (cm/s)
Time(min)
****
*
WT
TG
TdTomato L+R
tTA L+R
Mobile time
c
d
Time spent moving (s)
Immobile time
e
f
Time spent immobile (s)

#### Supplemental figure 6: The effect of eIF4E reduction in D1-SPNs on locomotion speed and duration.

a) Time-course of the mean velocity throughout the novelty condition experiment, with data binned in five-minute intervals. b) Overall mean velocity over the two-hour experimental period (genotype*RNAi interaction, 2-way ANOVA, $F(1, 33)=6.197$, $p=0.018$). c) Time-course of the cumulative time spent moving throughout the novelty condition experiment, with data binned in five-minute intervals. d) Total time spent moving over the two-hour experimental period. e) Time-course of the cumulative time spent immobile throughout the novelty condition experiment, with data binned in five-minute intervals. f) Total time spent immobile over the two-hour experimental period. Data are presented as mean ± SEM, with mean indicated by bar height or connecting line and SEM by error bars or lines. Dots represent individual values for each mouse. Behavioural experiments were performed using 9 TRE-GFP.shmiR-4E/WT TdTomato$^{L+R}$ and 10 TRE-GFP.shmiR-4E/TG TdTomato$^{L+R}$ mice, and 8 TRE-GFP.shmiR-4E/WT tTA$^{L+R}$ and 10 TRE-GFP.shmiR-4E/TG tTA$^{L+R}$ mice. Significance is denoted as, * $p<0.05$, ** $p<0.01$, *** $p<0.001$ and **** $p<0.0001$, calculated with Tukey's post-hoc multiple comparisons test. Data are presented as mean ± SEM, with mean indicated by bar height and SEM by error bars in bar graphs, and with mean indicated by the connecting line and SEM by error lines with shading in line graphs. Dots represent individual values for each mouse. For full details of statistical analysis including negative results, refer to supplemental table 12.

### Supplemental tables

#### Supplementary Table 1 related to Figure 1

| | WT | | | TG | | | | |
|---|---|---|---|---|---|---|---|---|
| Parameter | Mean | SEM | N (mice) | Mean | SEM | N (mice) | Test value and type | p-value |
| D1-Cre WT increase in activity with novelty | 31.33 | 7.723 | 6 | | | | t=4.057 1-sample t-test | ** **0.0098** |
| A2a-Cre WT increase in activity with novelty | 83.5 | 16.75 | 8 | | | | t=4.986 1-sample t-test | ** **0.0016** |
| D1-Cre increase in activity with novelty | 31.33 | 7.723 | 6 | 57.17 | 8.356 | 6 | t=2.27 Unpaired t-test | * **0.0465** |
| A2a-Cre increase in activity with novelty | 83.5 | 16.75 | 8 | 61.5 | 12.76 | 6 | t=0.9835 Unpaired t-test | ns 0.3448 |

#### Supplementary Table 2 related to Figure 2

| | WT | | | TG | | | | |
|---|---|---|---|---|---|---|---|---|
| Parameter | Mean +/- SEM | 25%, median, 75% | N (mice) n (cells) | Mean +/- SEM | 25%, median, 75% | N (mice) n (cells) | Test value and type | p-value |
| D1+ capacitance | 36.66 +/- 3.318 | 27.01, 32.30, 48.58 | 5, 17 | 32.28 +/- 2.340 | 25.99, 28.09, 43.28 | 5, 19 | t=1.098, unpaired t-test | ns 0.2801 |
| D1+ RMP | -77.75 +/- 1.274 | -82.20, -79.00, -73.60 | 5, 17 | -74.63 +/- 1.410 | -77.90, -75.60, -68.50 | 5, 19 | t=1.628, unpaired t-test | ns 0.1128 |
| D1+ Rm | 80.36 +/- 6.806 | 53.63, 78.91, 101.5 | 5, 17 | 103.8 +/- 7.637 | 77.57, 109.5, 117.9 | 5, 19 | t=2.267, unpaired t-test | * **0.0299** |
| D1+ AP duration | 0.8718 +/- 0.043 | 0.76, 0.82, 0.95 | 5, 17 | 0.9505 +/- 0.061 | 0.78, 0.87, 1.01 | 5, 19 | t=1.030, unpaired t-test | ns 0.3101 |
| D1+ AP peak | 93.61 +/- 3.051 | 84.66, 98.62, 104.2 | 5, 17 | 85.39 +/- 5.024 | 80.92, 93.05, 98.79 | 5, 19 | t=1.359, unpaired t-test | ns 0.1830 |
| D1+ rheobase | 373.4 +/- 25.27 | 277.5, 402.0, 459.5 | 5, 17 | 294.7 +/- 23.79 | 224.0, 311.0, 377.0 | 5, 19 | t=2.267, unpaired t-test | * **0.0298** |
| D1+ max firing frequency | 29.29 +/- 3.823 | 17.0, 26.0, 42.0 | 5, 17 | 44.42 +/- 3.883 | 34.0, 42.0, 56.0 | 5, 19 | t=2.7650, unpaired t-test | ** **0.0091** |
| **D1+ Current-Frequency Plots** | | | | | | | | |
| | WT | | | TG | | | | |
| Current | Mean | SEM | N (mice) n (cells) | Mean | SEM | N (mice) n (cells) | Test value and type | p-value |
| 0 | 0 | 0 | 5, 17 | 0 | 0 | 5, 19 | Two-way-RM-ANOVA Genotype: $F_{(1, 34)}$=5.449; Current step: $F_{(1.304, 44.33)}$ =79.32; Genotype x Current: $F_{(9, 306)}$=5.547 | Genotype: * **0.0256**; Current: **** **<0.0001**; Genotype x Current: **** **<0.0001** |
| 50 | 0.353 | 0.353 | 5, 17 | 0 | 0 | 5, 19 | | |
| 100 | 1.412 | 1.412 | 5, 17 | 2.632 | 1.475 | 5, 19 | | |
| 150 | 2.824 | 1.895 | 5, 17 | 6.632 | 2.808 | 5, 19 | | |
| 200 | 5.647 | 2.493 | 5, 17 | 12.316 | 3.834 | 5, 19 | | |
| 250 | 9.294 | 3.307 | 5, 17 | 19.474 | 4.382 | 5, 19 | | |
| 300 | 12.824 | 4.055 | 5, 17 | 26.947 | 4.65 | 5, 19 | | |
| 350 | 16.942 | 4.505 | 5, 17 | 33.895 | 4.341 | 5, 19 | | |
| 400 | 23.529 | 4.401 | 5, 17 | 39.789 | 4.092 | 5, 19 | | |
| 450 | 28.118 | 3.599 | 5, 17 | 43.895 | 3.88 | 5, 19 | | |
| 50 | 0.267 | 0.267 | 5, 15 | 0 | 0 | 5, 15 | | |
| 100 | 2.933 | 1.711 | 5, 15 | 2.8 | 1.93 | 5, 15 | | |
| 150 | 9.067 | 2.914 | 5, 15 | 8.533 | 3.107 | 5, 15 | | |
| 200 | 19.067 | 4.065 | 5, 15 | 18.933 | 4.0655 | 5, 15 | | |
| 250 | 28.8 | 4.424 | 5, 15 | 32 | 3.283 | 5, 15 | | |
| 300 | 39.2 | 4.079 | 5, 15 | 44.133 | 3.847 | 5, 15 | | |
| 350 | 50 | 3.641 | 5, 15 | 53.067 | 5.461 | 5, 15 | | |
| 400 | 58.133 | 5.841 | 5, 15 | 59.867 | 7.862 | 5, 15 | | |
| 450 | 64.667 | 7.797 | 5, 15 | 65.467 | 9.982 | 5, 15 | | |
| **D1+ Sholl Analysis** | | | | | | | | |
| | WT | | | TG | | | | |
| Distance | Mean | SEM | N (mice) n (cells) | Mean | SEM | N (mice) n (cells) | Test value and type | p-value |
| 10 | 4.333 | 0.373 | 4, 9 | 6.357 | 0.52 | 6, 14 | Two-way-RM-ANOVA Genotype: $F_{(1, 21)}$=0.3824; | Genotype: ns 0.543; |
| 30 | 10.778 | 1.234 | 4, 9 | 10.286 | 0.861 | 6, 14 | | |
| 50 | 11.111 | 0.992 | 4, 9 | 11.5 | 1.592 | 6, 14 | | |
| 70 | 9 | 0.972 | 4, 9 | 7.571 | 0.817 | 6, 14 | | |

| | | | | | | | | |
|---|---|---|---|---|---|---|---|---|
| 90 | 5.667 | 0.645 | 4, 9 | 4.643 | 0.487 | 6, 14 | Distance: $F_{(2.641, 55.47)}$ =81.36; Genotype x Distance: $F_{(8, 168)}$=1.208 | Distance: **** **<0.0001**; Genotype x Distance: ns 0.297 |
| 110 | 3.111 | 0.633 | 4, 9 | 1.929 | 0.425 | 6, 14 | | |
| 130 | 1.444 | 0.503 | 4, 9 | 0.429 | 0.251 | 6, 14 | | |
| 150 | 0.333 | 0.236 | 4, 9 | 0 | 0 | 6, 14 | | |
| 170 | 0.111 | 0.111 | 4, 9 | 0 | 0 | 6, 14 | | |

#### Supplementary Table 3 related to Figure 3

| | WT | | | TG | | | | |
|---|---|---|---|---|---|---|---|---|
| Parameter | Mean +/- SEM | 25%, median, 75% | N (mice) n (cells) | Mean +/- SEM | 25%, median, 75% | N (mice) n (cells) | Test value and type | p-value |
| D1+ P10 capacitance | 57.191 +/- 5.050 | 43.2, 52.5, 61.2 | 7, 23 | 56.925 +/- 5.164 | 45.625, 51.4, 64.55 | 5, 20 | Two-way-ANOVA Genotype: $F_{(1, 78)}$=1.418; Age: $F_{(1, 78)}$ =2.306; Genotype x Age: $F_{(1, 78)}$=1.302 | Genotype: ns 0.237; Age: ns 0.1329; Genotype x Age: ns 0.2573 |
| D1+ P28 capacitance | 71.457 +/- 5.110 | 53.2, 69.4, 88.3 | 5, 23 | 58.95 +/- 6.039 | 44.025, 54.7, 69.575 | 4, 16 | | |
| D1+ P10 RMP | -74.646 +/- 1.341 | -78.744, -76.385, -69.52 | 7, 23 | -80.679 +/- 1.312 | -84.896, -81.127, -77.278 | 5, 20 | Two-way-ANOVA Genotype: $F_{(1, 78)}$=0.419; Age: $F_{(1, 78)}$ =6.221; Genotype x Age: $F_{(1, 78)}$=22.26 | Genotype: ns 0.7074; Age: * **0.0147**; Genotype x Age: **** **<0.0001** |
| D1+ P28 RMP | -83.186 +/- 0.849 | -85.877, -83.328, -81.288 | 5, 23 | -78.045 +/- 1.051 | -81.32, -77.579, -75.637 | 4, 16 | | |
| D1+ P10 Rm | 167.649 +/- 8.36 | 139.6, 159.03, 191.679 | 7, 23 | 136.951 +/- 8.529 | 104.515, 125.85, 155.418 | 5, 20 | Two-way-ANOVA Genotype: $F_{(1, 78)}$=0.4113; Age: $F_{(1, 78)}$ =41.95; Genotype x Age: $F_{(1, 78)}$=7.49 | Genotype: ns 0.5232; Age: **** **<0.0001**; Genotype x Age: ** **0.0077** |
| D1+ P28 Rm | 83.919 +/- 9.616 | 49.097, 67.941, 101.685 | 5, 23 | 102.961 +/- 8.984 | 73.52, 102.32, 124.95 | 4, 16 | | |
| D1+ P10 AP duration | 1.118 +/- 0.071 | 0.845, 1.048, 1.284 | 7, 23 | 1.066 +/- 0.067 | 0.851, 1.023, 1.133 | 5, 20 | Two-way-ANOVA Genotype: $F_{(1, 78)}$=0.0025; Age: $F_{(1, 78)}$ =0.3298; Genotype x Age: $F_{(1, 78)}$=0.5124 | Genotype: ns 0.9606; Age: ns 0.5674; Genotype x Age: ns 0.4762 |
| D1+ P28 AP duration | 1.108 +/- 0.065 | 0.913, 1.006, 1.224 | 5, 23 | 1.153 +/- 0.052 | 0.981, 1.132, 1.238 | 4, 16 | | |
| D1+ P10 AP peak | 79.1 +/- 3.263 | 69.728, 81.467, 86.159 | 7, 23 | 84.628 +/- 3.172 | 79.344, 86.502, 90.883 | 5, 20 | Two-way-ANOVA Genotype: $F_{(1, 78)}$=0.0011; Age: $F_{(1, 78)}$ =0.3058; Genotype x Age: $F_{(1, 78)}$=2.836 | Genotype: ns 0.9173; Age: ns 0.5818; Genotype x Age: ns 0.0962 |
| D1+ P28 AP peak | 86.927 +/- 3.682 | 80.298, 94.672, 98.745 | 5, 23 | 80.671 +/- 3.581 | 76.181, 83.541, 88.765 | 4, 16 | | |
| D1+ P10 rheobase | 183.989 +/- 9.585 | 143.825, 192.803, 217.522 | 7, 21 | 195.852 +/- 6.25 | 179.41, 197.411, 211.801 | 5, 20 | Two-way-ANOVA Genotype: $F_{(1, 74)}$=0.7598; Age: $F_{(1, 74)}$ =69.9; Genotype x Age: $F_{(1, 74)}$=2.842 | Genotype: ns 0.3862; Age: **** **<0.0001**; |
| D1+ P28 rheobase | 330.359 +/- 18.044 | 266.246, 353.31, 395.202 | 5, 22 | 293.097 +/- 21.349 | 248.693, 283.047, 357.066 | 4, 15 | | |

| | | | | | | | | |
|---|---|---|---|---|---|---|---|---|
| | | | | | | | | Genotype x Age: ns 0.0960 |
| D1+ P10 max firing frequency | 57.217 +/- 2.847 | 44.0, 60.0, 70.0 | 7, 23 | 55.3 +/- 2.543 | 47.0, 56.0, 65.0 | 5, 20 | Two-way-ANOVA Genotype: $F_{(1, 78)}$=1.175; Age: $F_{(1, 78)}$ =56.39; Genotype x Age: $F_{(1, 78)}$=3.276 | Genotype: ns 0.2818; Age: **** **<0.0001**; Genotype x Age: ns 0.0741 |
| D1+ P28 max firing frequency | 32.609 +/- 2.072 | 26.0, 32.0, 40.0 | 5, 23 | 40.25 +/- 3.01 | 32.5, 41.0, 50.0 | 4, 16 | | |

**P10 Current-Frequency Plots**

| | WT | | | TG | | | | |
|---|---|---|---|---|---|---|---|---|
| Current | Mean | SEM | N (mice) n (cells) | Mean | SEM | N (mice) n (cells) | Test value and type | p-value |
| 0 | 0 | 0 | 7, 23 | 0 | 0 | 5, 20 | Two-way-RM-ANOVA Genotype: $F_{(1, 41)}$=1.333; Current step: $F_{(2.236, 91.68)}$ =194.4; Genotype x Current step: $F_{(9, 369)}$=0.181 | Genotype: ns 0.255; Current: **** **<0.0001**; Genotype x Current: ns 0.996 |
| 50 | 1.043 | 0.664 | 7, 23 | 0 | 0 | 5, 20 | | |
| 100 | 7.652 | 2.248 | 7, 23 | 4 | 1.2482 | 5, 20 | | |
| 150 | 21.652 | 2.819 | 7, 23 | 17.9 | 2.15 | 5, 20 | | |
| 200 | 34.696 | 2.258 | 7, 23 | 30.9 | 2.054 | 5, 20 | | |
| 250 | 43.913 | 1.99 | 7, 23 | 40.1 | 1.732 | 5, 20 | | |
| 300 | 49.652 | 2.0378 | 7, 23 | 46.4 | 2.106 | 5, 20 | | |
| 350 | 51.217 | 2.696 | 7, 23 | 49.1 | 2.911 | 5, 20 | | |
| 400 | 52.087 | 4.198 | 7, 23 | 50 | 3.503 | 5, 20 | | |
| 450 | 51.13 | 4.657 | 7, 23 | 48.9 | 4.546 | 5, 20 | | |

**P28 Current-Frequency Plots**

| | WT | | | TG | | | | |
|---|---|---|---|---|---|---|---|---|
| Current | Mean | SEM | N (mice) n (cells) | Mean | SEM | N (mice) n (cells) | Test value and type | p-value |
| 0 | 0 | 0 | 5, 23 | 0 | 0 | 4, 16 | Two-way-RM-ANOVA Genotype: $F_{(1, 37)}$=2.833; Current step: $F_{(1.269, 46.96)}$ =143.7; Genotype x Current step: $F_{(9, 333)}$=2.635 | Genotype: ns 0.1008; Current: **** **<0.0001**; Genotype x Current: ** **0.0059** |
| 50 | 0 | 0 | 5, 23 | 0 | 0 | 4, 16 | | |
| 100 | 0.348 | 0.348 | 5, 23 | 0.875 | 0.657 | 4, 16 | | |
| 150 | 2.087 | 1.235 | 5, 23 | 3.5 | 2.094 | 4, 16 | | |
| 200 | 4.087 | 2.022 | 5, 23 | 7.125 | 3.093 | 4, 16 | | |
| 250 | 8.087 | 2.513 | 5, 23 | 14.5 | 3.819 | 4, 16 | | |
| 300 | 13.478 | 2.976 | 5, 23 | 22.25 | 4.074 | 4, 16 | | |
| 350 | 20.348 | 2.833 | 5, 23 | 28.75 | 4.037 | 4, 16 | | |
| 400 | 27.304 | 2.488 | 5, 23 | 35 | 3.54 | 4, 16 | | |
| 450 | 32.609 | 2.0716 | 5, 23 | 40 | 2.921 | 4, 16 | | |

#### Supplementary Table 4 related to Figure 4

| | WT | | | TG | | | | |
|---|---|---|---|---|---|---|---|---|
| Parameter | Mean +/- SEM | 25%, median, 75% | N (mice) n (cells) | Mean +/- SEM | 25%, median, 75% | N (mice) n (cells) | Test value and type | p-value |
| TdTomato capacitance | 69.602 +/- 5.688 | 53.736, 63.841, 87.179 | 4, 14 | 58.475 +/- 3.72 | 46.424, 60.734, 67.401 | 4, 12 | Two-way-ANOVA Genotype: $F_{(1, 54)}$=1.083; RNAi: $F_{(1, 54)}$ =0.0049; | Genotype: ns 0.3026; RNAi: ns 0.9445; Genotype x RNAi: ns 0.2318 |
| tTA capacitance | 63.276 +/- 3.525 | 51.681, 61.696, 74.311 | 4, 17 | 64.109 +/- 6.076 | 48.088, 55.923, 68.112 | 4, 15 | | |

| | | | | | | | | |
|---|---|---|---|---|---|---|---|---|
| | | | | | | | Genotype x RNAi: $F_{(1, 54)}$=1.462 | |
| TdTomato RMP | -74.579 +/- 1.571 | -81.525, -73.9, -69.875 | 4, 14 | -73.233 +/- 0.88 | -75.9, -73.0, -71.45 | 4, 12 | Two-way-ANOVA Genotype: $F_{(1, 54)}$=0.4618; RNAi: $F_{(1, 54)}$ =0.5316; Genotype x RNAi: $F_{(1, 54)}$=2.556 | Genotype: ns 0.4997; RNAi: ns 0.4691; Genotype x RNAi: ns 0.1157 |
| tTA RMP | -73.306 +/- 1.231 | -77.95, -73.2, -69.0 | 4, 17 | -76.64 +/- 1.809 | -80.2, -77.2, -72.0 | 4, 15 | | |
| TdTomato Rm | 90.451 +/- 9.435 | 61.643, 87.725, 101.268 | 4, 14 | 99.066 +/- 5.66 | 85.004, 94.9, 114.761 | 4, 12 | Two-way-ANOVA Genotype: $F_{(1, 54)}$=0.0029; RNAi: $F_{(1, 54)}$ =0.6120; Genotype x RNAi: $F_{(1, 54)}$=1.534 | Genotype: ns 0.9572; RNAi: ns 0.4375; Genotype x RNAi: ns 0.2209 |
| tTA Rm | 105.147 +/- 6.744 | 93.782, 102.357, 115.304 | 4, 17 | 95.749 +/- 6.159 | 76.114, 93.957, 108.6 | 4, 15 | | |
| TdTomato rheobase | 261.629 +/- 30.133 | 150.425, 274.750, 374.7 | 4, 14 | 208.608 +/- 13.784 | 180.850, 212.850, 246.1 | 4, 12 | Two-way-ANOVA Genotype: $F_{(1, 53)}$=0.0458; RNAi: $F_{(1, 53)}$ =0.7079; Genotype x RNAi: $F_{(1, 53)}$=8.028 | Genotype: ns 0.8314; RNAi: ns 0.4039; Genotype x RNAi: ** **0.0065** |
| tTA rheobase | 187.247 +/- 12.103 | 152.6, 183.9, 231.8 | 4, 17 | 248.929 +/- 20.244 | 167.05, 265.6, 290.925 | 4, 14 | | |
| TdTomato max firing frequency | 32.714 +/- 2.378 | 25.5, 32.0, 41.0 | 4, 14 | 41.5 +/- 2.935 | 34.0, 40.0, 50.0 | 4, 12 | Two-way-ANOVA Genotype: $F_{(1, 54)}$=0.074; RNAi: $F_{(1, 54)}$ =12.20; Genotype x RNAi: $F_{(1, 54)}$=10.10 | Genotype: ns 0.7865; RNAi: *** **0.001**; Genotype x RNAi: ** **0.0024** |
| tTA max firing frequency | 31.765 +/- 3.262 | 22.0, 36.0, 42.0 | 4, 17 | 21.333 +/- 3.072 | 12.0, 18.0, 30.0 | 4, 15 | | |
| **TdTomato Current-Frequency Plots** | | | | | | | | |
| | WT | | | TG | | | | |
| Current | Mean | SEM | N (mice) n (cells) | Mean | SEM | N (mice) n (cells) | Test value and type | p-value |
| 0 | 0 | 0 | 4, 14 | 0 | 0 | 4, 12 | Three-way-RM-ANOVA Genotype: $F_{(1, 54)}$=0.6179; Current: $F_{(2.12, 114.5)}$ =94.54; RNAi: $F_{(1, 54)}$ =3.53; Genotype x Current: $F_{(9, 486)}$=1.944; RNAi x Current: $F_{(9, 486)}$=6.041; Genotype x RNAi: $F_{(1, 54)}$=12.04; | Genotype: ns 0.4353; Current: **** **<0.0001**; RNAi: ns 0.0657; Genotype x Current: * **0.044**; Genotype x RNAi: ** **0.001**; Current x RNAi: **** **<0.0001**; |
| 50 | 0.143 | 0.143 | 4, 14 | 0 | 0 | 4, 12 | | |
| 100 | 2.857 | 1.952 | 4, 14 | 0 | 0 | 4, 12 | | |
| 150 | 6.286 | 2.904 | 4, 14 | 1.167 | 0.796 | 4, 12 | | |
| 200 | 10.714 | 3.707 | 4, 14 | 9.5 | 2.676 | 4, 12 | | |
| 250 | 12.571 | 4.18 | 4, 14 | 19 | 3.521 | 4, 12 | | |
| 300 | 16.571 | 4.319 | 4, 14 | 27.167 | 3.655 | 4, 12 | | |
| 350 | 20.286 | 4.027 | 4, 14 | 33.167 | 3.68 | 4, 12 | | |
| 400 | 24.714 | 3.417 | 4, 14 | 38.333 | 3.103 | 4, 12 | | |
| 450 | 28.143 | 3.448 | 4, 14 | 41.5 | 2.935 | 4, 12 | | |
| **tTA Current-Frequency Plots** | | | | | | | | |
| | WT | | | TG | | | | |
| Current | Mean | SEM | N (mice) n (cells) | Mean | SEM | N (mice) n (cells) | | |
| 0 | 0 | 0 | 4, 17 | 0 | 0 | 4, 15 | | |
| 50 | 0.235 | 0.235 | 4, 17 | 0 | 0 | 4, 15 | | |

| | | | | | | | | |
|---|---|---|---|---|---|---|---|---|
| 100 | 2.588 | 1.739 | 4, 17 | 0.933 | 0.547 | 4, 15 | Genotype x Current x RNAi: $F_{(9, 486)}$=6.46; | Genotype x Current x RNAi: **** **<0.0001** |
| 150 | 9.176 | 2.432 | 4, 17 | 2.933 | 1.498 | 4, 15 | | |
| 200 | 15.765 | 2.975 | 4, 17 | 5.6 | 1.914 | 4, 15 | | |
| 250 | 22.471 | 2.872 | 4, 17 | 7.067 | 2.046 | 4, 15 | | |
| 300 | 25.059 | 3.005 | 4, 17 | 11.6 | 2.546 | 4, 15 | | |
| 350 | 26.118 | 3.664 | 4, 17 | 13.467 | 2.777 | 4, 15 | | |
| 400 | 25.294 | 3.861 | 4, 17 | 16.267 | 3.23 | 4, 15 | | |
| 450 | 23.765 | 4.246 | 4, 17 | 17.2 | 3.743 | 4, 15 | | |

#### Supplementary Table 5 related to Figure 5

| | WT | | | TG | | | | |
|---|---|---|---|---|---|---|---|---|
| Parameter | Mean +/- SEM | 25%, median, 75% | N (mice) n (cells) | Mean +/- SEM | 25%, median, 75% | N (mice) n (cells) | Test value and type | p-value |
| CsCl2 D1-TdTomato sensitivity | 41.25 +/- 6.482 | 15.54, 44.19, 55.65 | 6, 11 | 35.87 +/- 5.564 | 30.53 32.41 46.14 | 4, 7 | t=0.5779, unpaired t-test | ns 0.5714 |
| XE991 D1-TdTomato sensitivity | 29.168 +/- 2.857 | 20.056, 28.841, 39.029 | 5, 18 | 15.385 +/- 3.523 | 4.098, 15.134, 24.777 | 5, 11 | Two-way-ANOVA Genotype: $F_{(1, 51)}$=3.823; RNAi: $F_{(1, 51)}$ =0.0902; Genotype x RNAi: $F_{(1, 51)}$=5.218 | Genotype: ns 0.056; RNAi: ns 0.7652; Genotype x RNAi: * **0.0266** |
| XE991 tTA sensitivity | 22.718 +/- 4.059 | 17.394, 20.515, 31.753 | 4,10 | 23.787 +/- 2.608 | 14.476, 20.156, 32.485 | 6, 16 | | |

#### Supplementary Table 6 related to Figure 6

| | WT | | | TG | | | | |
|---|---|---|---|---|---|---|---|---|
| Parameter | Mean | SEM | N (mice) | Mean | SEM | N (mice) | Test value and type | p-value |
| TdTomato Total distance travelled | 14149.7 | 713.89 | 9 | 24761.9 | 1696.9 | 10 | Two-way-ANOVA Genotype: $F_{(1, 33)}$=22.37; RNAi: $F_{(1, 33)}$ =1.982; Genotype x RNAi: $F_{(1, 33)}$=7.607 | Genotype: **** **<0.0001**; RNAi: ns 0.1685; Genotype x RNAi: ** **0.0094** |
| tTA Total distance travelled | 16063.4 | 1572.16 | 8 | 18857.7 | 1389.41 | 10 | | |
| TdTomato Total marbles buried | 5.333 | 0.85 | 9 | 11.5 | 1.628 | 10 | Two-way-ANOVA Genotype: $F_{(1, 33)}$=1.288; RNAi: $F_{(1, 33)}$ =2.159; Genotype x RNAi: $F_{(1, 33)}$=13.13 | Genotype: ns 0.2645; RNAi: ns 0.1512; Genotype x RNAi: *** **0.001** |
| tTA Total marbles buried | 8.125 | 1.674 | 8 | 4.9 | 0.823 | 10 | | |

#### Supplementary Table 7 related to Supplementary Figure 1

| | WT | | | TG | | | | |
|---|---|---|---|---|---|---|---|---|
| Parameter | Mean | SEM | N (mice) | Mean | SEM | N (mice) | Test value and type | p-value |
| D1-Cre frequency | 43.5 | 13.98 | 6 | 91.5 | 12.27 | 6 | t=2.581 Unpaired t-test | * **0.0274** |
| D1-Cre amplitude | 46.67 | 11.73 | 6 | 58.83 | 9.336 | 6 | t=0.8115 Unpaired t-test | ns 0.4360 |
| D1-Cre AUC | 50.33 | 15.38 | 6 | 117.2 | 25.52 | 6 | t=2.243 Unpaired t-test | * **0.0487** |
| A2a-Cre frequency | 98.38 | 20.65 | 8 | 55.17 | 13.71 | 6 | t=1.613 Unpaired t-test | ns 0.1327 |
| A2a-Cre amplitude | 110.3 | 24.47 | 8 | 98.0 | 21.73 | 6 | t=0.3598 Unpaired t-test | ns 0.7252 |
| A2a-Cre AUC | 127.5 | 29.14 | 8 | 89.17 | 22.79 | 6 | t=0.9786 Unpaired t-test | ns 0.3471 |

#### Supplementary Table 8 related to Supplementary Figure 2

| | Baseline | | | SCH39166 | | | | |
|---|---|---|---|---|---|---|---|---|
| Parameter | Mean | SEM | N (mice) | Mean | SEM | N (mice) | Test value and type | p-value |
| Frequency | 0.225 | 0.0275 | 6 | 0.06933 | 0.0133 | 6 | t=5.062 Paired t-test | ** **0.0039** |
| Amplitude | 1.217 | 0.1898 | 6 | 0.4333 | 0.0707 | 6 | t=4.875 Paired t-test | ** **0.0046** |
| AUC | 35172 | 3996 | 6 | 16735 | 4510 | 6 | t=5.554 Paired t-test | ** **0.0026** |

#### Supplementary Table 9 related to Supplementary Figure 3

| | WT | | | TG | | | | |
|---|---|---|---|---|---|---|---|---|
| Parameter | Mean +/- SEM | 25%, median, 75% | N (mice) n (cells) | Mean +/- SEM | 25%, median, 75% | N (mice) n (cells) | Test value and type | p-value |
| D1-capacitance | 33.43 +/- 2.596 | 24.70, 33.50, 42.65 | 5, 15 | 24.44 +/- 3.502 | 14.91, 18.31, 26.51 | 5, 15 | t=2.062, unpaired t-test | * **0.0486** |
| D1- RMP | -76.31 +/- 1.325 | -79.40, -75.10, -72.70 | 5, 15 | -74.98 +/- 0.792 | -77.90, -75.10, -73.00 | 5, 15 | t=0.8593, unpaired t-test | ns 0.3975 |
| D1- Rm | 103.1 +/- 9.420 | 78.44, 91.59, 118.0 | 5, 15 | 128.1 +/- 11.42 | 93.34, 118.2, 177.0 | 5, 15 | t=1.694, unpaired t-test | ns 0.1014 |
| D1- AP duration | 0.832 +/- 0.0824 | 0.71, 0.78, 0.94 | 5, 15 | 0.9767 +/- 0.0768 | 0.79, 0.90, 1.25 | 5, 15 | t=1.284, unpaired t-test | ns 0.2095 |
| D1- AP peak | 86.28 +/- 5.374 | 65.62, 95.04, 101.7 | 5, 15 | 77.74 +/- 4.82 | 63.45, 85.48, 92.51 | 5, 15 | t=1.184, unpaired t-test | ns 0.2464 |

| | | | | | | | | |
|---|---|---|---|---|---|---|---|---|
| D1- rheobase | 239.1 +/- 18.84 | 177.0, 242.0, 296.0 | 5, 15 | 247.2 +/- 13.62 | 198.0, 259.0, 289.0 | 5, 15 | t=0.3470, unpaired t-test | ns 0.7312 |
| D1- max firing frequency | 65.73 +/- 7.424 | 46.0, 56.0, 68.0 | 5, 15 | 65.73 +/- 9.877 | 44.0, 52.0, 62.0 | 5, 15 | t=0.0000, unpaired t-test | ns >0.999 |
| **D1- Current-Frequency Plots** | | | | | | | | |
| | WT | | | TG | | | | |
| Current | Mean | SEM | N (mice) n (cells) | Mean | SEM | N (mice) n (cells) | Test value and type | p-value |
| 0 | 0 | 0 | 5, 15 | 0 | 0 | 5, 15 | Two-way-RM-ANOVA Genotype: $F_{(1, 28)}$=0.1375; Current step: $F_{(1.937, 54.25)}$ =75.69; Genotype x Current step: $F_{(9, 252)}$=0.1057 | Genotype: ns 0.7136; Current: **** **<0.0001**; Genotype x Current: ns 0.9995 |
| 50 | 0.267 | 0.267 | 5, 15 | 0 | 0 | 5, 15 | | |
| 100 | 2.933 | 1.711 | 5, 15 | 2.8 | 1.93 | 5, 15 | | |
| 150 | 9.067 | 2.914 | 5, 15 | 8.533 | 3.107 | 5, 15 | | |
| 200 | 19.067 | 4.065 | 5, 15 | 18.933 | 4.0655 | 5, 15 | | |
| 250 | 28.8 | 4.424 | 5, 15 | 32 | 3.283 | 5, 15 | | |
| 300 | 39.2 | 4.079 | 5, 15 | 44.133 | 3.847 | 5, 15 | | |
| 350 | 50 | 3.641 | 5, 15 | 53.067 | 5.461 | 5, 15 | | |
| 400 | 58.133 | 5.841 | 5, 15 | 59.867 | 7.862 | 5, 15 | | |
| 450 | 64.667 | 7.797 | 5, 15 | 65.467 | 9.982 | 5, 15 | | |
| **D1- Sholl Analysis** | | | | | | | | |
| | WT | | | TG | | | | |
| Distance | Mean | SEM | N (mice) n (cells) | Mean | SEM | N (mice) n (cells) | Test value and type | p-value |
| 10 | 4.571 | 0.481 | 4, 7 | 6 | 1.517 | 4, 5 | Two-way-RM-ANOVA Genotype: $F_{(1, 10)}$=0.014; Distance: $F_{(2.259, 22.59)}$ =35.09; Genotype x Distance: $F_{(8, 80)}$=1.195 | Genotype: ns 0.9087; Distance: **** **<0.0001**; Genotype x Distance: ns 0.3126 |
| 30 | 8.571 | 1.131 | 4, 7 | 11.6 | 1.4 | 4, 5 | | |
| 50 | 10.429 | 2.057 | 4, 7 | 12 | 0.707 | 4, 5 | | |
| 70 | 8.571 | 1.757 | 4, 7 | 8.2 | 1.356 | 4, 5 | | |
| 90 | 6.429 | 2.125 | 4, 7 | 4.4 | 1.03 | 4, 5 | | |
| 110 | 2.857 | 0.67 | 4, 7 | 1.6 | 0.678 | 4, 5 | | |
| 130 | 1 | 0.436 | 4, 7 | 0.4 | 0.4 | 4, 5 | | |
| 150 | 0.714 | 0.286 | 4, 7 | 0 | 0 | 4, 5 | | |
| 170 | 0 | 0 | 4, 7 | 0 | 0 | 4, 5 | | |

#### Supplementary Table 10 related to Supplementary Figure 4

| | WT | | | TG | | | | |
|---|---|---|---|---|---|---|---|---|
| Parameter | Mean +/- SEM | 25%, median, 75% | N (mice) n (cells) | Mean +/- SEM | 25%, median, 75% | N (mice) n (cells) | Test value and type | p-value |
| D1- P10 capacitance | 57.361 +/- 4.659 | 42.8, 55.1, 66.6 | 7, 23 | 45.848 +/- 3.116 | 34.9, 46.2, 54.0 | 5, 23 | Two-way-ANOVA Genotype: $F_{(1, 80)}$=3.365; Age: $F_{(1, 80)}$ =4.908; Genotype x Age: $F_{(1, 80)}$=0.1321 | Genotype: ns 0.0703; Age: * **0.0296**; Genotype x Age: ns 0.7172 |
| D1- P28 capacitance | 67.063 +/- 6.715 | 50.0, 60.2, 84.6 | 5, 19 | 59.358 +/- 6.481 | 35.9, 56.6, 79.8 | 4, 19 | | |
| D1- P10 RMP | -74.867 +/- 1.527 | -81.863, -75.165, -70.287 | 7, 23 | -74.627 +/- 1.668 | -80.442, -73.0, -69.06 | 5, 23 | Two-way-ANOVA Genotype: $F_{(1, 80)}$=0.0243; Age: $F_{(1, 80)}$ =18.37; Genotype x Age: $F_{(1, 80)}$=0.0001 | Genotype: ns 0.8766; Age: **** **<0.0001**; |
| D1- P28 RMP | -81.029 +/- 1.109 | -85.2, -81.325, -77.375 | 5, 19 | -80.82 +/- 1.149 | -83.72, -82.069, -76.915 | 4, 19 | | |

| | | | | | | | | |
|---|---|---|---|---|---|---|---|---|
| | | | | | | | | Genotype x Age: ns 0.9914 |
| D1- P10 Rm | 204.675 +/- 20.995 | 149.4, 177.84, 220.66 | 7, 23 | 225.667 +/- 22.039 | 159.91, 190.58, 255.9 | 5, 23 | Two-way-ANOVA Genotype: $F_{(1, 80)}$=0.5317; Age: $F_{(1, 80)}$ =46.78; Genotype x Age: $F_{(1, 80)}$=0.2018 | Genotype: ns 0.468; Age: **** **<0.0001**; Genotype x Age: ns 0.6545 |
| D1- P28 Rm | 90.779 +/- 9.275 | 63.48, 73.55, 114.25 | 5, 19 | 95.768 +/- 9.45 | 60.33, 83.08, 123.8 | 4, 19 | | |
| D1- P10 AP duration | 1.189 +/- 0.054 | 1.0, 1.157, 1.481 | 7, 23 | 1.191 +/- 0.073 | 0.933, 1.196, 1.286 | 5, 23 | Two-way-ANOVA Genotype: $F_{(1, 80)}$=0.7236; Age: $F_{(1, 80)}$ =0.0233; Genotype x Age: $F_{(1, 80)}$=0.6628 | Genotype: ns 0.3975; Age: ns 0.8790; Genotype x Age: ns 0.4180 |
| D1- P28 AP duration | 1.146 +/- 0.06 | 0.936, 1.069, 1.406 | 5, 19 | 1.254 +/- 0.07 | 1.07, 1.306, 1.435 | 4, 19 | | |
| D1- P10 AP peak | 73.01 +/- 2.634 | 65.992, 72.393, 82.441 | 7, 23 | 71.956 +/- 4.342 | 52.947, 72.095, 90.182 | 5, 23 | Two-way-ANOVA Genotype: $F_{(1, 80)}$=0.5; Age: $F_{(1, 80)}$ =5.179; Genotype x Age: $F_{(1, 80)}$=0.1979 | Genotype: ns 0.4816; Age: * **0.0255**; Genotype x Age: ns 0.6576 |
| D1- P28 AP peak | 83.94 +/- 4.357 | 68.997, 80.987, 102.004 | 5, 19 | 79.312 +/- 4.629 | 70.0, 82.474, 96.717 | 4, 19 | | |
| D1- P10 rheobase | 162.724 +/- 10.511 | 123.7, 150.736, 192.002 | 7, 23 | 150.647 +/- 11.051 | 112.575, 139.718, 184.189 | 5, 23 | Two-way-ANOVA Genotype: $F_{(1, 78)}$=0.4480; Age: $F_{(1, 78)}$ =96.68; Genotype x Age: $F_{(1, 78)}$=2.285 | Genotype: ns 0.5053; Age: **** **<0.0001**; Genotype x Age: ns 0.1347 |
| D1- P28 rheobase | 282.046 +/- 18.126 | 221.249, 295.868, 352.158 | 5, 19 | 313.319 +/- 18.518 | 255.854, 334.63, 365.329 | 4, 17 | | |
| D1- P10 max firing frequency | 51.826 +/- 2.238 | 44.0, 50.0, 60.0 | 7, 23 | 56.87 +/- 3.689 | 50.0, 54.0, 68.0 | 5, 23 | Two-way-ANOVA Genotype: $F_{(1, 80)}$=0.8777; Age: $F_{(1, 80)}$ =17.11; Genotype x Age: $F_{(1, 80)}$=0.5773 | Genotype: ns 0.3517; Age: **** **<0.0001**; Genotype x Age: ns 0.4496 |
| D1- P28 max firing frequency | 41.789 +/- 2.324 | 34.0, 42.0, 52.0 | 5, 19 | 42.316 +/- 3.172 | 36.0, 42.0, 50.0 | 4, 19 | | |
| **P10 Current-Frequency Plots** | | | | | | | | |
| | WT | | | TG | | | | |
| Current | Mean | SEM | N (mice) n (cells) | Mean | SEM | N (mice) n (cells) | Test value and type | p-value |
| 0 | 0 | 0 | 7, 23 | 0 | 0 | 5, 23 | Two-way-RM-ANOVA Genotype: $F_{(1, 44)}$=0.6895; Current step: $F_{(2.286, 100.6)}$ =54.49; Genotype x Current step: $F_{(9, 396)}$=0.9744 | Genotype: ns 0.4108; Current: **** **<0.0001**; Genotype x Current: ns 0.4606 |
| 50 | 4.087 | 1.613 | 7, 23 | 0.957 | 0.485 | 5, 23 | | |
| 100 | 17.913 | 3.037 | 7, 23 | 15.826 | 2.589 | 5, 23 | | |
| 150 | 30.087 | 3.323 | 7, 23 | 29.13 | 3.723 | 5, 23 | | |
| 200 | 37.652 | 3.19 | 7, 23 | 38.348 | 3.788 | 5, 23 | | |
| 250 | 41.304 | 2.591 | 7, 23 | 43.217 | 3.966 | 5, 23 | | |
| 300 | 39.478 | 3.0314 | 7, 23 | 45.043 | 4.491 | 5, 23 | | |
| 350 | 37.217 | 3.863 | 7, 23 | 45.652 | 5.119 | 5, 23 | | |
| 400 | 35.739 | 4.59 | 7, 23 | 43.913 | 5.88 | 5, 23 | | |
| 450 | 34.087 | 5.118 | 7, 23 | 41.652 | 6.158 | 5, 23 | | |
| **P28 Current-Frequency Plots** | | | | | | | | |
| | WT | | | TG | | | | |
| Current | Mean | SEM | N (mice) n (cells) | Mean | SEM | N (mice) n (cells) | Test value and type | p-value |
| 0 | 0 | 0 | 5, 19 | 0 | 0 | 4, 19 | | Genotype: |

| | | | | | | | | |
|---|---|---|---|---|---|---|---|---|
| 50 | 0 | 0 | 5, 19 | 0 | 0 | 4, 19 | Two-way-RM-ANOVA Genotype: $F_{(1, 36)}=0.1098$; Current step: $F_{(1.46, 52.57)}=131.0$; Genotype x Current step: $F_{(9, 324)}=0.1398$ | ns 0.7423; Current: **** **<0.0001**; Genotype x Current: ns 0.9985 |
| 100 | 1.158 | 0.799 | 5, 19 | 0.526 | 0.428 | 4, 19 | | |
| 150 | 3.895 | 1.967 | 5, 19 | 4 | 1.892 | 4, 19 | | |
| 200 | 8.842 | 2.923 | 5, 19 | 9.053 | 3.195 | 4, 19 | | |
| 250 | 16.526 | 3.5961 | 5, 19 | 13.895 | 4.117 | 4, 19 | | |
| 300 | 24.316 | 3.63 | 5, 19 | 22.105 | 4.223 | 4, 19 | | |
| 350 | 31.895 | 3.166 | 5, 19 | 30.316 | 3.882 | 4, 19 | | |
| 400 | 37.579 | 2.616 | 5, 19 | 36.105 | 3.659 | 4, 19 | | |
| 450 | 41.263 | 2.254 | 5, 19 | 39.894 | 3.76 | 4, 19 | | |

#### Supplementary Table 11 related to Supplementary Figure 5

| | TdTomato | | | tTA | | | | |
|---|---|---|---|---|---|---|---|---|
| Parameter | Mean | SEM | N | Mean | SEM | N | Test value and type | p-value |
| eIF4E cell density | 736.6 | 119.7 | 4 slices, 3 mice | 151.4 | 71.08 | 5 slices, 3 mice | t=4.417 Unpaired t-test | ** **0.0031** |
| WT western blot | 100 | 7.676 | 3 | 43.081 | 13.423 | 3 | Two-way-ANOVA Genotype: $F_{(1, 8)}=3.608$; RNAi: $F_{(1, 8)}=61.21$; Genotype x RNAi: $F_{(1, 8)}=5.836$ | Genotype: ns 0.0941; RNAi: **** **<0.0001**; Genotype x RNAi: * **0.0421** |
| TG western blot | 145.418 | 13.906 | 3 | 37.645 | 3.263 | 3 | | |

#### Supplementary Table 12 related to Supplementary Figure 6

| | WT | | | TG | | | | |
|---|---|---|---|---|---|---|---|---|
| Parameter | Mean | SEM | N (mice) | Mean | SEM | N (mice) | Test value and type | p-value |
| TdTomato Mean velocity | 1.998 | 0.1 | 9 | 3.485 | 0.238 | 10 | Two-way-ANOVA Genotype: $F_{(1, 33)}=23.82$; RNAi: $F_{(1, 33)}=1.613$; Genotype x RNAi: $F_{(1, 33)}=6.197$ | Genotype: **** **<0.0001**; RNAi: ns 0.2129; Genotype x RNAi: * **0.018** |
| TTA Mean velocity | 2.244 | 0.219 | 8 | 2.726 | 0.206 | 10 | | |
| TdTomato Time spent mobile | 2231.4 | 125.29 | 9 | 3314.5 | 189.91 | 10 | Two-way-ANOVA Genotype: $F_{(1, 33)}=15.21$; RNAi: $F_{(1, 33)}=0.8886$; Genotype x RNAi: $F_{(1, 33)}=2.526$ | Genotype: *** 0.0004; RNAi: ns 0.3527; Genotype x RNAi: ns 0.1215 |
| tTA Time spent mobile | 2359.0 | 229.18 | 8 | 2814.8 | 222.8 | 10 | | |
| TdTomato Time spent immobile | 4871.7 | 149.36 | 9 | 3813.5 | 187.7 | 10 | Two-way-ANOVA Genotype: $F_{(1, 33)}=18.31$; RNAi: $F_{(1, 33)}=0.436$; Genotype x RNAi: $F_{(1, 33)}=0.9768$ | Genotype: *** **0.0002**; RNAi: ns 0.5137; Genotype x RNAi: ns 0.3302 |
| TTA Time spent immobile | 4805.8 | 223.94 | 8 | 4144.7 | 226.77 | 10 | | |
